# Selection, Design and Immunogenicity Studies of ASFV Antigens for Subunit mRNA Cocktail Vaccines with Specific Immune Response Profiles

**DOI:** 10.1101/2024.10.08.617156

**Authors:** Fangfeng Yuan, Junru Cui, Tianlei Wang, Jane Qin, Ju Hyeong Jeon, Huiming Ding, Charles A. Whittaker, Renhuan Xu, Helen Cao, Jianzhu Chen

## Abstract

Development of safe and effective subunit vaccines for controlling African Swine Fever Virus (ASFV) infection has been hampered by a lack of protective viral antigens, complex virion structures, and multiple mechanisms of infection. Here, we selected ASFV antigens based on their localization on the virion, known functions, and homologies to the subunits of the protective vaccinia virus vaccine. We also engineered viral capsid proteins for inducing optimal antibody responses and designed T cell-directed antigen for inducing broad and robust cellular immunity. The selected antigens in lipid nanoparticle-mRNA formulations were evaluated for immunogenicity in both mice and pigs with concordant results. Different antigens induced divergent immune response profiles, including the levels of IgG and T cell responses and effector functions of anti-sera. We further developed a computational approach to combine antigens into cocktails for inducing specific immune response profiles and validated candidate cocktail vaccines in mice. Our results provide a basis for further evaluating candidate subunit mRNA vaccines in challenge studies.

**Teaser:** Novel strategies to develop subunit vaccines for ASFV and other complex large DNA viruses.

## Introduction

African swine fever (ASF) is a highly contagious swine viral disease that causes almost 100% mortality in domestic pigs. Currently the United States is free of ASF and incursion to the US swine population could result in approximately $50 billion losses (*1*). The causative agent, African swine fever virus (ASFV), is a large DNA virus with a double-stranded DNA genome of 170 to 190 kb, encoding at least 150 proteins (*2*). Although a live attenuated vaccine was licensed in Vietnam in 2023, it has major safety concerns (*3*). A safe and effective subunit ASF vaccine is urgently needed.

ASF vaccine development faces several major hurdles (*4*): i) ASFV is classified as a biosafety level 3 (BSL-3) agent. The requirement for BSL-3 containment for any virus work limits the research on the virus and vaccine development. ii) Viral antigens to elicit protective immunity against ASFV infection have not been identified (*4*), impeding the development of effective subunit vaccines. iii) ASFV exhibits complex multi-layer structures. Both the extracellular enveloped virion (EEV), i.e., with the outer membrane, and intracellular mature virion (IMV), i.e., without the outer membrane, are infectious (*5*). iv) ASFV can infect host macrophages through multiple entry mechanisms, including receptor-mediated endocytosis, clathrin-mediated endocytosis, macropiniocytosis, and phagocytosis (*6*). As a result, there is no reliable viral neutralization assay for measuring anti-viral antibody responses induced by any single antigen (*7*). v) Immunological correlates of protective immunity against ASFV are not fully defined. Studies have shown that antibody responses elicited by inactivated virus were not sufficient to protect pigs from infection (*8*), whereas immune responses by live attenuated virus confer protection (*9*). Therefore, T cell immunity likely play a pivotal role in the protection. Lymphocyte depletion studies showed that cytotoxic CD8^+^ lymphocytes are important for ASFV clearance (*10*). IFN-γ responses correlate with the degree of cross-protection against heterologous ASFV challenge (*11*). Therefore, an effective ASFV vaccine must induce both humoral and cellular immunities.

ASFV is the only member in *Asfarviridae*, which belongs to the phylum of nucleocytoplasmic large DNA viruses (NCLDV), characterized by the complex virion structures, large genomes and protein-coding capacity (*2*). *Poxviridae*, a well-characterized family in NCLDV, is the most closely related to *Asfarviridae* phylogenetically (**Figure S1**). As a key member of *Poxviridae*, the use of vaccinia virus (VACV) as a vaccine has contributed tremendously to the eradication of smallpox disease caused by Variola virus (*12*). Like ASFV, VACV also produces two infectious virions: EEV and IMV. Studies have shown that subunit vaccines, composing two EEV antigens (B5R and A33R) and two IMV antigens (L1R and A27L), provide complete protection against lethal VACV challenge in mice (*13–16*), suggesting the feasibility of developing safe and effective subunit vaccines for NCLDV with appropriate viral immunogens.

The success of lipid nanoparticle (LNP)-delivered mRNA encoding the spike protein as COVID-19 vaccines motivates broad applications of mRNA-based vaccines. In mRNA vaccines, proteins are synthesized by the host cells and likely maintain native structural conformation, and therefore could induce both humoral and cellular immunities. Antibodies produced by B cells could block viral infection of host cells through neutralization and help to inactivate or eliminate viruses through other effector mechanisms, such as antibody-dependent complement deposition (ADCP), antibody-dependent cellular phagocytosis (ADCP), and antibody-dependent cellular cytotoxicity (ADCC). T cells could recognize virus-infected cells and help to clear the virus through cytokines or directly killing of the infected cells. In addition, mRNA vaccines are especially suited for developing subunit vaccines as multiple mRNAs can be formulated into the same LNP, greatly simplifying the manufacturing process.

In this study, we rationally selected and comprehensively evaluated immunogenicity of ASFV antigens, including i) ASFV homologs corresponding to the subunits of the protective VACV vaccine; ii) promising immunogens reported in previous studies; iii) viral capsid proteins in membrane-bound form for more efficient induction of antibody responses; and iv) novel multiple T cell epitopes (MTE) for inducing broad and robust cellular immunity. In mRNA-LNP formulation, the selected ASFV antigens stimulated concordant antigen-specific antibody and T cell responses in both mice and pigs, with the MTE inducing the most potent cellular immunity. By combining the effector function profiles of antigen-specific antibodies and the levels of B and T cell responses induced by different ASFV antigens, we developed optimal antigen combinations for cocktail vaccines with specific immune response profiles, which were further validated in mice. Our study represents a comprehensive investigation of ASF mNRA subunit vaccines incorporating rational antigen selection, protein engineering, T cell-directed antigen design, and profiling of immune responses. Our results provide a basis for further evaluating the subunit mRNA vaccine candidates in challenge studies. The innovative strategies reported here should be applicable to design vaccines for other large complex DNA viruses such as monkeypox.

## Results

### Selection and design of ASFV candidate antigens

To identify viral antigens for a safe and effective subunit mRNA vaccine for ASF, we selected ASFV antigens using the following approaches. First, we selected ASFV homologs corresponding to the subunits of the protective VACV vaccine because ASFV is most closely related to VACV (**Figure S1**) and both viruses produce two types of infectious virions (EEV and IMV). Four VACV antigens were used in the subunit vaccine, including A33R and B5R from EEV and L1R and A27L from IMV (**Table 1**). Thus, we selected ASFV EP153R, encoding C-type lectin, which is localized on the outer membrane and shares 18% amino acid identity with VACV A33R and ASFV CD2-like protein (CD2v or EP402R), a major outer membrane protein with a size around 41 kDa, which shares 12% amino acid identity with VACV B5R (42 kDa) (**Figure 1A**). L1R, localized on IMV of VACV, is responsible for membrane fusion. In ASFV, both E248R and E199L have been reported to play a similar role (*6, 17*) and share 14% and 8% amino acid identities with L1R, respectively. VACV A27L exists as a trimer in the capsid on IMV and contributes to attachment to cell heparan sulfate receptor (*18, 19*). In ASFV, the viral capsid is formed by two proteins, the trimeric P72 and pentameric Penton (*20*). Because both P72 and Penton are intracellular proteins and exist in multimeric forms in the capsid, we have engineered them as membrane-bound (MB) form to preserve the multimeric structures and to more effectively induce antibody responses (*21*). In addition, a single mutation (N180Q) was introduced into the MB-Penton to abolish the glycosylation. Therefore, MB-P72 and MB-P_N180Q_ are included here as comparison. For simplicity, MB-P72 and MB-P_N180Q_ are referred to as P72 and Penton in this study.

**Fig. 1.**
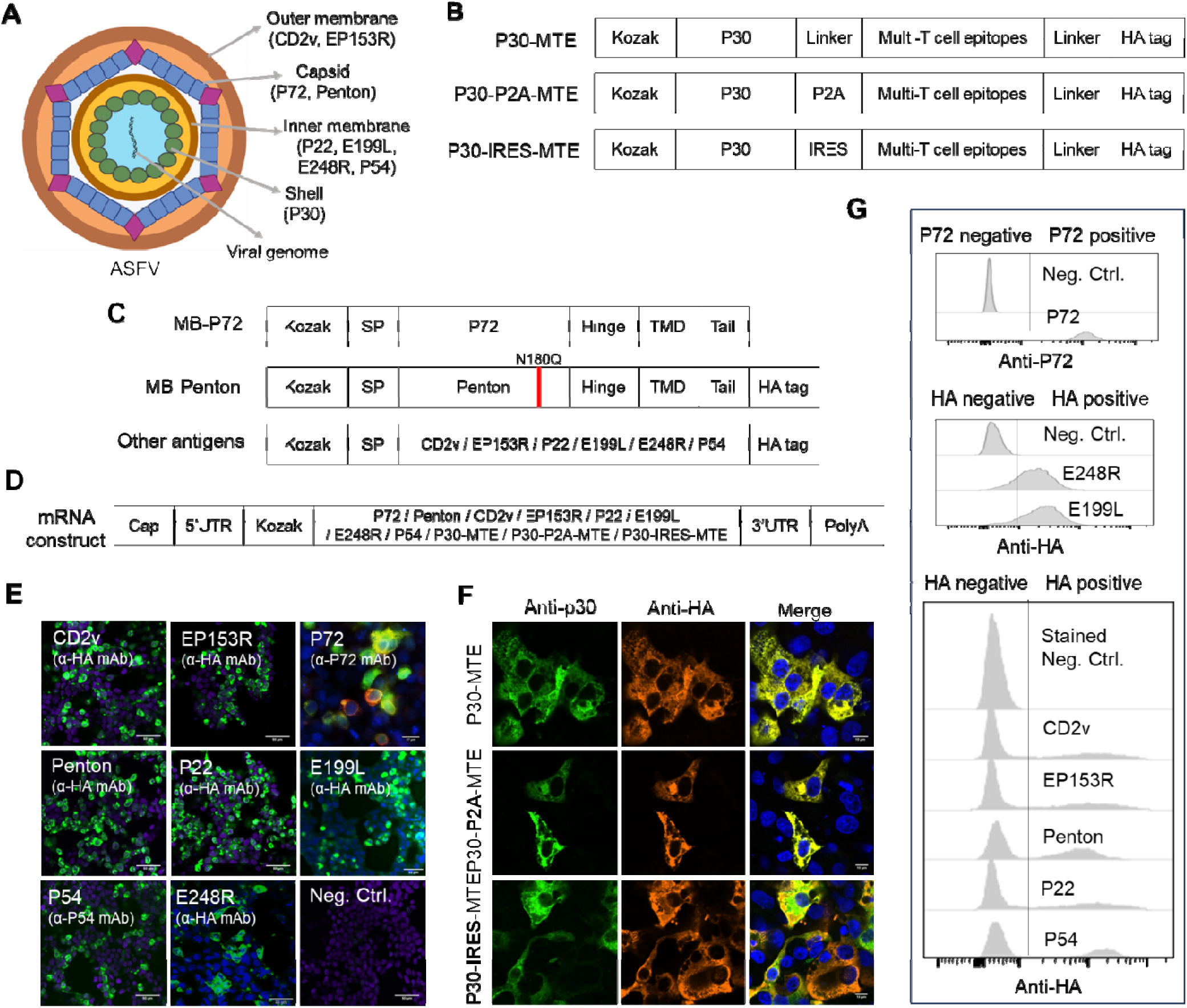
Selection, design, and in vitro validation of ASFV vaccine candidate antigens. (**A**) schematic diagram of the extracellular enveloped form of ASFV. Each layer of the viral particle is shown as distinctive colors, including the outer membrane, capsid layer, inner membrane, core shell, and viral DNA genome. Antigens selected in this study are depicted under each layer. (**B**) Schematic diagrams showing the three different designs for expressing T cell-directed antigens. In P30-MTE, P30 is fused with multi-T cell epitope (MTE) by a GGGS linker. In P30-P2A-MTE, P30 is linked with MTE via P2A self-cleavage site. In P30-IRES-MTE, MTE is translated separately through IRES. All three constructs contain an HA tag in the C terminus for detecting MTE expression. Kozak sequence was added in the N terminus of all constructs in aid of translation. (**C**) Schematic diagrams of DNA expression vectors for P72, Penton, CD2v, EP153R, P22, E199L, E248R, and P54. The membrane-anchoring of P72 and Penton was accomplished by addition of signal peptide (SP) in the N terminus, a hinge, transmembrane domain (TMD) and short cytoplasmic tail in the C terminus. The red vertical line represents N180Q mutation to remove the glycan on Penton. A HA tag was added in the C terminus of all other ASFV proteins except P72 for easy monitoring expression at the protein level. For P72 detection, a commercial monoclonal antibody was used. (**D**) Schematic diagrams of mRNA construct design. All designed ASFV genes were inserted into a pUC vector between the 5’ untranslated region (5’ UTR) and the 3’UTR, followed by polyA. A cap was added to the in vitro transcribed mRNA using Vaccinia Capping Enzyme. For testing immunogenicity in mice, all mRNA constructs, except P72, included a HA tag in the C terminus. The HA tag was removed for immunogenicity testing in pigs. (**E-F**) Confocal imaging of candidate antigens in HEK 293T cells transfected with DNA plasmids expressing CD2v, EP153R, P72, Pen.ton, P22, E199L, E248R, and P54 (**E**), or P30-MTE, P30-P2A-MTE, and P30-IRES-MTE (**F**). Scale bar is 50 μm. (**G**) Flow cytometry analysis for expression of ASFV antigens following lipid nanoparticle (LNP)-mRNA transfection of HEK 293T cells. For confocal microscopy and flow cytometry, monolayer HEK 293T cells were transfected with DNA plasmids by polyethylenimine or LNP-mRNAs. Cells were harvested and fixed at 48 hours post transfection and stained with mouse mAbs specific for P72, P54, P30, and mouse anti-HA tag mAb for CD2v, EP153R, P22, E199L, E248R. A rabbit anti-HA tag polyclonal antibody was used to stain MTE antigens. AF488-conjugated goat anti-mouse IgG was used as secondary antibody and an additional AF594 conjugated goat-anti-rabbit IgG was used for **F**. Nuclei were stained with DAPI before confocal imaging.

**Table 1.** Comparison of vaccinia virus protective antigens with ASFV homologs.

| VACV <sup>1</sup> |  | ASFV |  | AA identity | Localization | Shared feature |
| --- | --- | --- | --- | --- | --- | --- |
| Name | Size | Name | Size |  |  |  |
| A33R | 23-28 kDa | EP153R | 18 kDa | 18% | EEV <sup>2</sup> | C-type lectin-like |
| B5R | 42 kDa | CD2v | 41 kDa | 12% | EEV | Outer membrane |
| L1R | 27 kDa | E248R | 27 kDa | 14% | IMV <sup>3</sup> | Fusion protein |
|  |  | E199L | 24 kDa | 8% |  |  |
| A27L | 14 kDa | P72 | 73 kDa | N/A <sup>*</sup> | IMV | Trimer |
|  |  | Penton | 28 kDa |  |  | Pentamer |

<sup>1</sup> Vaccinia virus (VACV)
<sup>2</sup> Extracellular enveloped virus (EEV)
<sup>3</sup> Intracellular mature virion (IMV)
\* N/A: not applicable

Second, we selected ASFV antigens that have shown to be promising. ASFV P72 and CD2v were reported to induce neutralizing antibodies, which provide partial protection against lethal viral challenge (*22–28*). ASFV EP153R was shown to contain several T cell epitopes and synergized with CD2v in reducing viremia and disease symptoms (*29, 30*). ASFV P54 was shown to induce neutralizing antibody response or partial protection against viral challenge (*8, 31*). ASFV P22 is a highly immunogenic protein with application for serological diagnostics and potential role in maintaining virion structure (*32, 33*). Hence, besides P72, CD2v and EP153R, P54 and P22, both localized on the viral membrane, were also selected as candidate antigens for evaluation (**Figure 1A**).

Third, we developed a novel multi-T cell epitope (MTE) antigen to stimulate strong cellular immunity against ASFV. T cell epitopes were selected based on i) searching IEDB database and literature for experimentally identified epitopes by IFN-γ ELISpot and/or MHC/mass spectrometry using recovered pig lymphocytes, and ii) NetMHCpan prediction of epitopes that bind to the most frequently swine leukocyte antigen (SLA) alleles (SLA-1:0101, SLA-1:0401 and SLA-1:0801) and those with highest binding affinity from abundantly expressed viral proteins (*34, 35*) were selected. A total of 27 epitopes were selected (**Table 2**), including 22 experimentally identified epitopes (6 from EP153R, 4 from PP62, 5 from MGF100-1L, 1 each from A238L, K145R, MGF505-7R, P34, and P37, 2 from P150) (*10, 30, 36–40*), and 5 predicted epitopes (3 from M448R and 2 from MGF505-7R). Notably, the 6 known T cell epitopes from CD2v and 1 from P72 were not included in the MTE design as these two antigens were tested separately. All epitopes were fused together with additional 5 endogenous amino acid residues at each end for proteasomal degradation and in an order that avoid disordered polypeptide structure based on Alphafold 2 prediction. In addition, to facilitate the expression of the MTE antigen, the highly expressed ASFV P30 is linked to MTE by a direct GGGS linker (P30-MTE), or a P2A self-cleavage site (P30-P2A-MTE), or an internal ribosome entry site (IRES) signal (P30-IRES-MTE) for direct translation of MTE (**Figure 1B**). Notably, P30 is highly immunogenic and has also been reported to induce neutralizing antibodies (*8, 31, 41*).

**Table 2.**
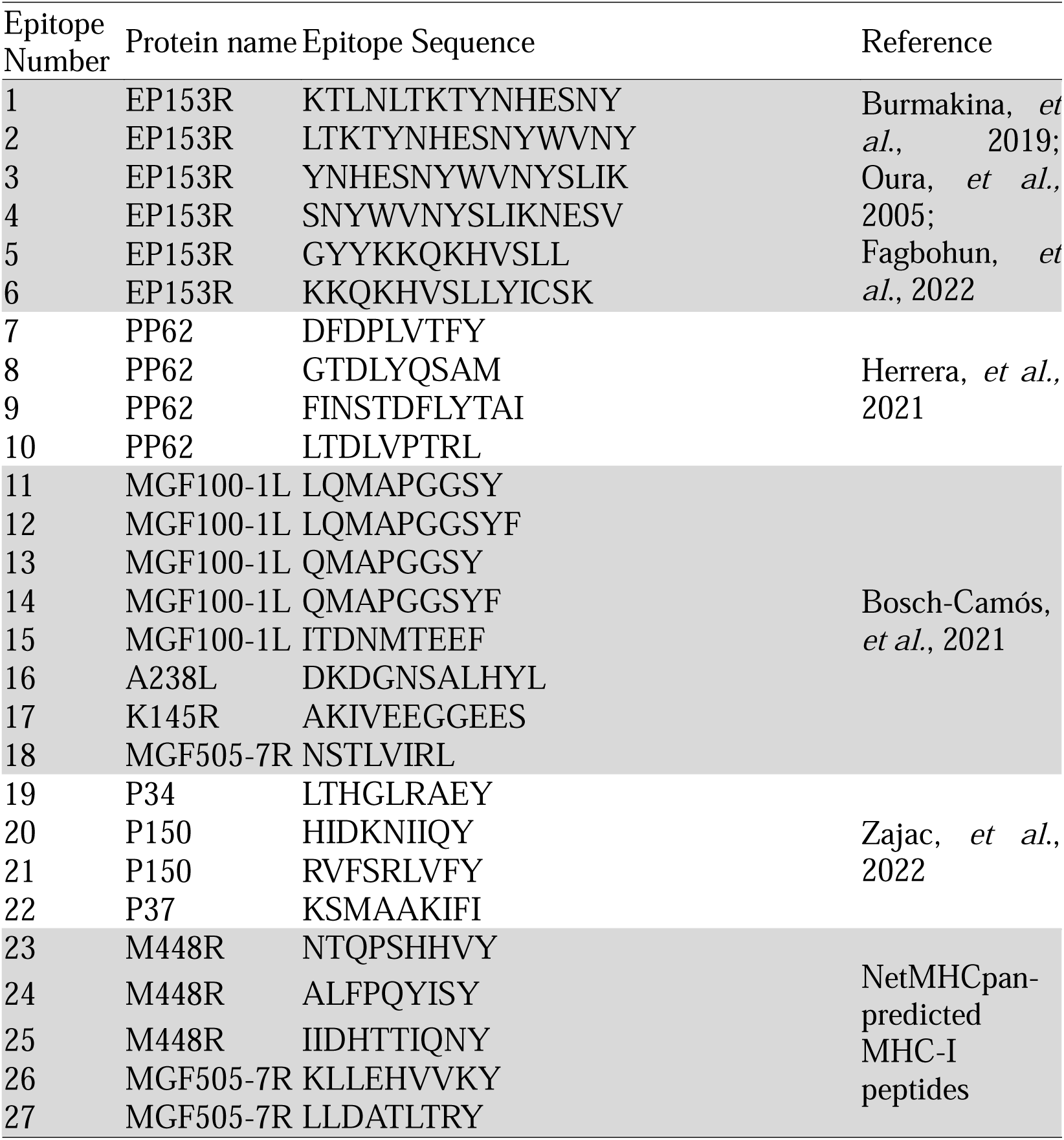
T cell epitope sequences and their viral protein origins in MTE.

In total, we constructed and tested 11 vectors expressing CD2v, EP153R, P72, Penton, P22, E199L, E248R, P54, P30-MTE, P30-P2A-MTE, and P30-IRES-MTE (**Figure 1A-C** and **Figure S2A**). Because specific antibodies are not available for all selected ASFV proteins, except P72, P54 and P30, we added a HA tag at the C-terminus of the most proteins for easy monitoring of their expression. The HA tag was removed for immunogenicity studies in pigs.

### Validation of expression of candidate antigens *in vitro*

We validated expression of the selected ASFV candidate antigens in cell lines. HEK 293T and/or Vero cells were transfected with individual expression vectors (**Figure S2A**) followed by confocal immunofluorescence imaging with antibodies specific for P72, P54, and P30, and anti-HA antibody for the rest. Abundant fluorescence signals were detected for all vectors in the transfected but not untransfected cells (**Figure 1E**). Expressions of CD2v, P22, P54, EP153R, E199L, and E248R were also detected by Western blotting (**Figure S3**). Notably, CD2v and EP153R were highly glycosylated, consistent with previous reports (*42, 43*). Expression of P72 and Penton were validated previously (*21*). To validate co-expression of P30 and MTE, transfected HEK 293T cells were co-stained with anti-P30 and anti-HA antibodies, followed by confocal microscopy. Both P30 and MTE from all three different designs were readily detected in the cytosol of transfected cells (**Figure 1F**). Thus, the selected ASFV antigens can be expressed in cell lines.

Next, we prepared mRNA for each candidate antigen by in vitro transcription and formulated the mRNA in LNP. Briefly, DNA fragments encoding P72, Penton, CD2v, EP153R, E199L, E248R, P22, P54, P30-MTE, P30-P2A-MTE, and P30-IRES-MTE were inserted into a pUC plasmid containing the T7 promoter, 5’UTR, 3’UTR, and polyA (**Figure 1D** and **Figure S2B**). Plasmids were linearized and used as templates for *in vitro* transcription using T7 RNA polymerase and ATP, GTP, CTP and pseudo-UTP. mRNAs were capped, purified, and formulated into LNPs using ionizable lipid, DSPC, cholesterol and DMG-PEG-2000 as described (*44*). The LNP-mRNA formulations were further subject to physicochemical analysis and results showed an average particle size of 80 – 120 nm in diameter and a low poly dispersity index (PDI) of 0.1 to 0.2 (**Table S1**), suggesting that the LNP-mRNA particles have a uniform particle size distribution, optimal for particle internalization and biodistribution. LNP-mRNAs were transfected into HEK 293T cells and expression of all candidate antigens was readily detected by flow cytometry (**Figure 1G**). These results suggest that formulated LNP-mRNAs are efficiently translated.

### Induction of antibody and T cell responses in mice by LNP-mRNA vaccination

We first evaluated the immunogenicity of the candidate antigens in mice. BALB/c mice at 6-8 weeks of age were divided into 12 groups with 5 mice per group. Eleven groups were immunized with 11 LNP-mRNAs expressing different antigens twice at day 0 and day 21 (**Figure 2A**). The other group was injected with sterile PBS and served as control. Blood was collected before immunization and at day 14 and 35 after immunization for assessing antigen-specific IgG responses by ELISA. Spleen was collected at day 35 for assessing antigen-specific T cell responses by ELISPOT. Compared to before immunization, antigen-specific IgG responses, as indicated by the endpoint titers, were significantly induced in all immunized mice 14 days after the first immunization and further boosted at day 35 (**Figure 2B**). However, the levels of IgG titers varied significantly among the different antigens. For example, 14 days after the first immunization antigen-specific IgG titer was only 1.2-fold over the background for E248R whereas the titer was 125.2-fold for P22. Fourteen days following the boost, antigen-specific IgG titer was increased to 126 (2.2-fold) for E248R and to 521,675 (83-fold) for P22. Among the 11 different antigens, P22, P30-IRES-MTE and P30-P2A-MTE induced the highest titers of antigen-specific IgG responses after boost, followed by E199L, P72, Penton, P54 and CD2v, with the lowest for EP153R, E248R and P30-MTE (**Figure 2B**). Similarly, LNP-mRNA immunization induced significant antigen-specific T cell responses as indicated by secretion of cytokines IFN-γ and TFN-α following re-stimulation of splenocytes with either specific recombinant proteins or P72 peptides (**Figure 2C-D**). T cell responses also varied significantly among different antigens. For example, CD2v stimulated highest level of cytokine-secreting spots (IFN-γ: 327, TNF-α: 992), followed by EP153R (IFN-γ: 116, TNF-α: 414), P54 (IFN-γ: 59, TNF-α: 346), Penton (IFN-γ: 54, TNF-α: 593), P72 (IFN-γ: 30, TNF-α: 216), and P22 (IFN-γ: 29, TNF-α: 223). These results show that mRNA expressing the selected ASFV antigens induce both antibody and T cell responses, but the magnitudes of the immune responses vary considerably among the different antigens.

**Fig. 2.**
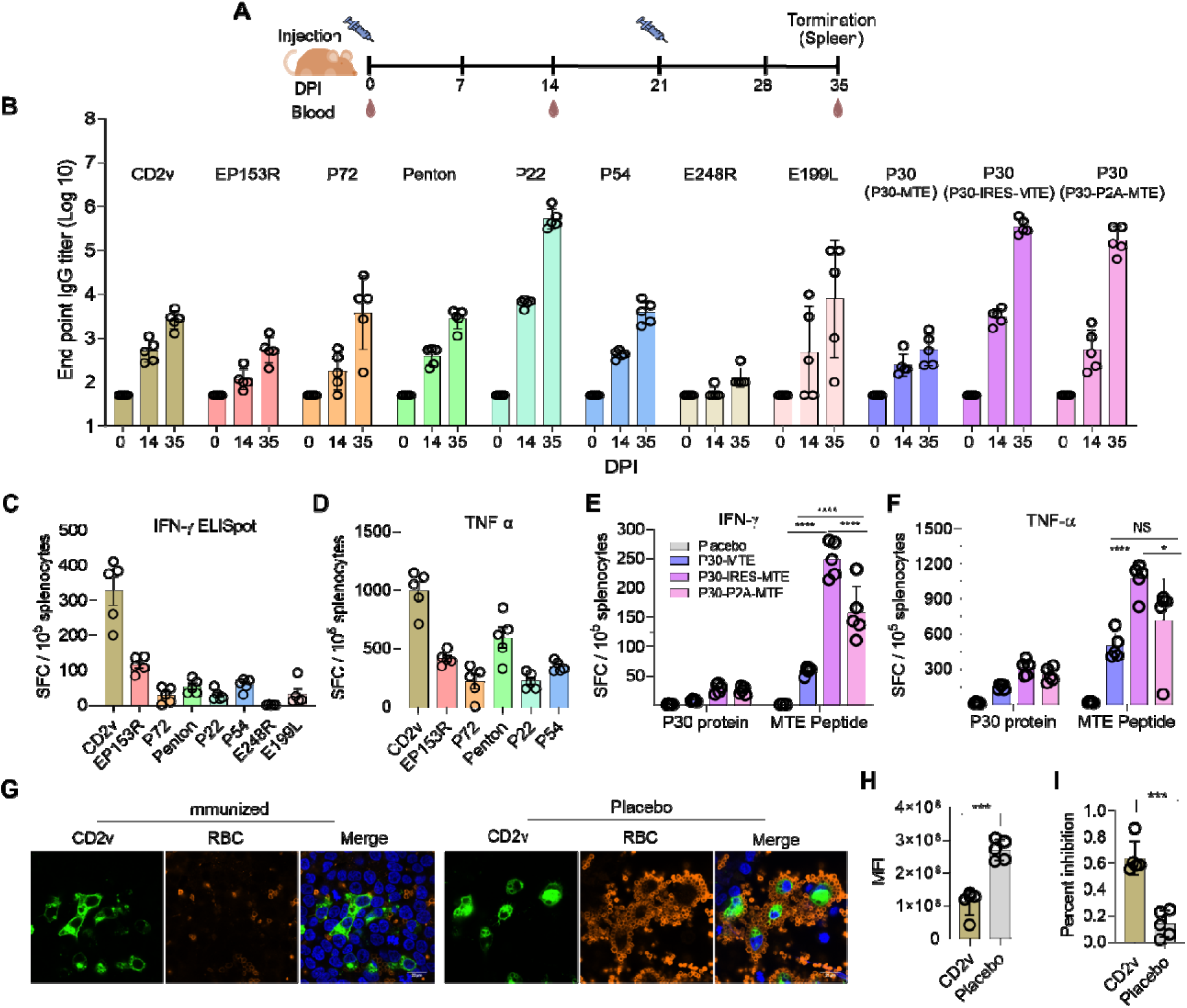
Immunogenicity of candidate antigens in mice. **(A)** Experimental design. Mice (5 per group) were immunized with 5 µg mRNA in LNP formulation at day 0 and day 21 or injected with PBS as control. Blood was collected before immunization and at 14 and 35 days post immunization (DPI) and used for assaying antigen-specific IgG levels by ELISA. Mice were euthanized at day 35 (14 days after boost) and spleen was collected for assaying antigen-specific T cell responses by ELISPOT. **(B)** Antigen-specific IgG responses normalized to day 0 (before immunization). For MTE groups, IgG titers specific for P30 were measured. The endpoint titer was defined as the reciprocal of the highest serum dilution that gives a reading above the cutoff which was determined by average OD_405_ readings of placebo mice + 3 standard deviation. **(C-D)** Antigen-specific T cell responses to CD2v, EP153R, P72, Penton, P22, P54, E248R and E199L. 10^5^ splenocytes from each mouse were seeded on ELISpot plate and stimulated with 5 µg/mL of purified proteins or P72 peptide. PMA/ionomycin cocktail and medium were included as positive and negative stimulation control, respectively. Dual-cytokine ELISpot was performed in 36 hours post stimulation. The number of IFN-γ (**C**) and TNF-α (**D**) spot-forming cells (SFC) per 10^5^ splenocytes was calculated and plotted. **(E-F)** Antigen-specific T cell responses to P30 (**E**) and MTE (**F**). ELISpot assay was done as in **C-D** except P30 protein and MTE peptide cocktail were used to stimulated splenocytes. **(G-I)** Hemoadsorption inhibition assay (HADIA) of CD2v antisera. Sera (n=5) from CD2v LNP-mRNA immunized mice were treated with receptor destroying enzyme (RDE) to remove unspecific factors inhibiting hemoadsorption, followed by incubation with CD2v-HA-expressing HEK 293T cells. 2% porcine RBC were added to the mixture and incubated for another 24 hours. Cells were fixed, permeabilized, and stained with a mouse anti-HA monoclonal antibody and rabbit anti-RBC polyclonal antibodies. AF594-labled anti-rabbit and AF488-labled anti-mouse IgG were added for secondary antibody staining. Cells were counterstained with DAPI before imaging using confocal microscope. Representative confocal images (**G**) and mean fluorescent intensity (**H**) and calculated percentage of inhibition (**I**). Each circle represents one mouse in B-F and H-I. One-way ANOVA for ELISpot assay and student’s t-test for HADIA were used for statistical analysis. NS, no significance; *, P<0.05; **, P<0.01; ***, P<0.001; ****, P<0.0001.

The three MTE designs also induced significantly different immune responses. P30-IRES-MTE induced the highest titer of anti-P30 IgG (**Figure 2B**) and highest numbers of MTE-specific IFN-γ and TNF-α spots (**Figure 2E-F**), followed by P30-P2A-MTE. P30-MTE induced the lowest antibody and T cell responses. Notably, the MTE-specific T cell responses, as indicated by the numbers of IFN-γ and TNF-α ELISpots, were significantly higher than P30-specific responses, indicating that the T cell-directed antigen (MTE) is potent. Compared to P30-MTE, where P30 is fused with MTE via a GGGS linker, in P30-P2A-MTE, P30-MTE is synthesized as a single polypeptide that is cleaved at P2A site, and in P30-IRES-MTE, P30 and MTE are synthesized separately through the internal ribosomal entry site (IRES). Our results show that a complete separation of P30 and MTE translation through IRES is most effective for inducing both P30-specific antibody responses and MTE-specific T cell responses.

ASFV induced hemadsorption is characterized by the adherence of red blood cells to the surface of infected cells (*45*). This phenomenon is primarily mediated by viral protein CD2v expressed on the surface of infected cells. To test if CD2v-specific antibodies inhibit hemadsorption, sera from CD2v LNP-mRNA immunized mice were first incubated with CD2v-expressing HEK 293T cells and then pig red blood cells (RBC) were added for surface adherence. Hemadsorption was observed by confocal microscopy after anti-RBC antibody staining. As shown in **Figure 2G**, sera from unimmunized mice did not show significant inhibition of hemadsorption or “rosette” formation surrounding CD2v-expressing cells, while the sera from CD2v LNP-mRNA immunized mice inhibited “rosette” formation. Quantification of percentages of inhibition showed that CD2v immunized sera yielded 64% inhibition of hemadsorption as compared to 13% with control sera (**Figure 2H-I**). These results show that although CD2v-specific IgG titer is relatively low compared to those induced by other ASFV antigens, the antibodies can specifically block CD2v-mediated hemadsorption.

### Induction of antibody and T cell responses in pigs by LNP-mRNA vaccination

We next evaluated immunogenicity of the following ASFV antigens in pigs based on their induction of relatively higher levels of antibody and T cell responses in mice, including P72, Penton, P22, E199L, P54, CD2v, EP153R, P30-IRES-MTE, P30-P2A-MTE. Commercial piglets at 6 weeks of age were immunized with 30 μg mRNA in LNP formulation (4 pigs per group) twice at day 0 and day 21 (**Figure 3A**). Two pigs were injected with sterile PBS and served as control. Blood was collected before immunization and every 7 days after immunization for assessing antigen-specific IgG responses by ELISA. Spleen were collected at day 35 for assessing antigen-specific T cell responses by ELISPOT. No clinical signs were observed throughout the study and immunized pigs gained as much weight as control pigs, suggesting that LNP-mRNA is safe for use in pigs.

**Fig. 3.**
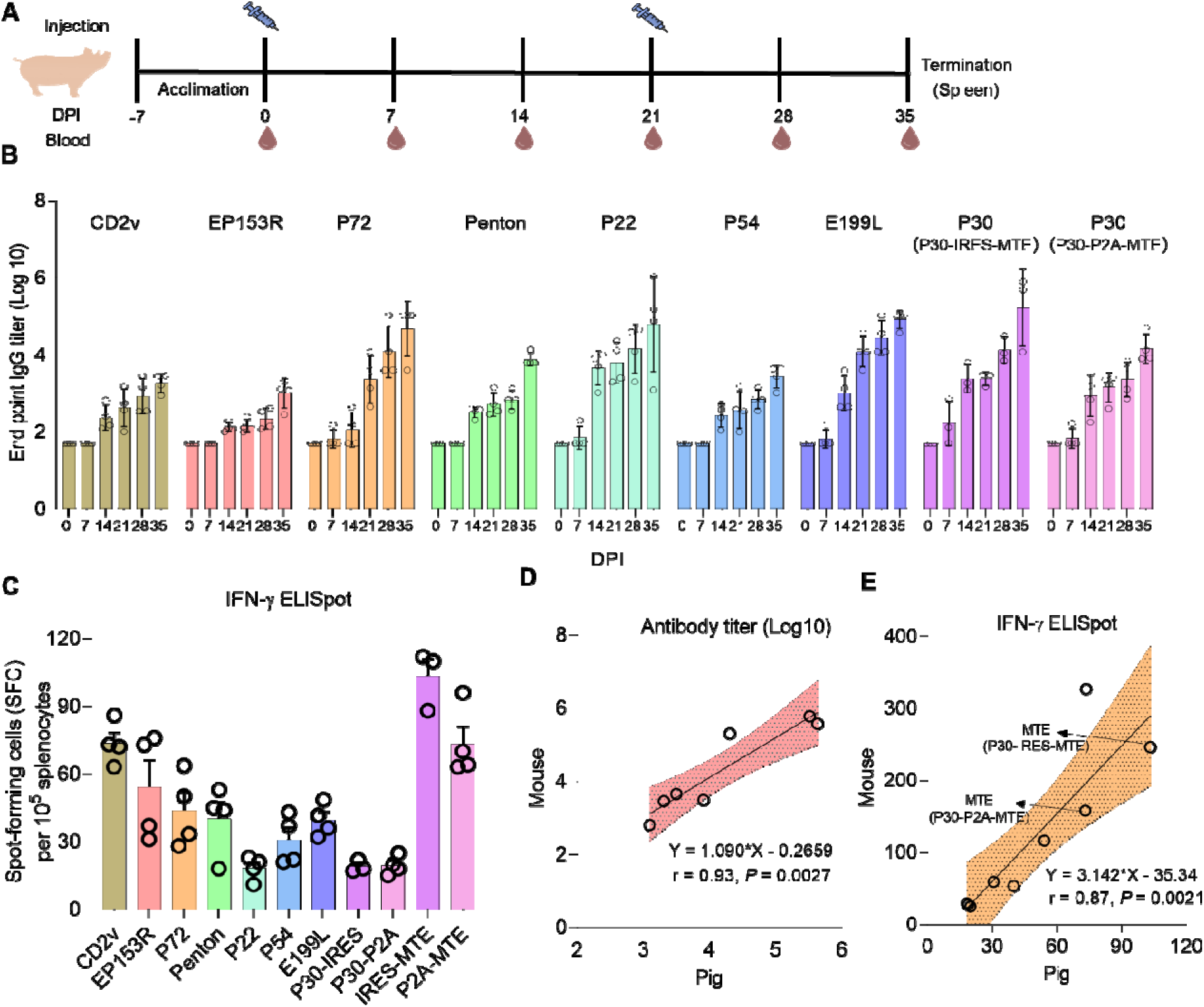
Immunogenicity of candidate antigens in pigs. **(A)** Experimental design. Piglets (4 per group) were immunized with 30 µg mRNA in LNP formulation at day 0 and day 21 or injected with PBS as control (2 pigs). Blood was collected before immunization and every 7 days post immunization (DPI) and used for assaying antigen-specific IgG levels by ELISA. Pigs were euthanized at day 35 (14 days after boost) and spleen was collected for assaying antigen-specific T cell responses by ELISPOT. **(B)** Antigen-specific IgG responses normalized to day 0 (before immunization). For MTE groups, IgG titers specific for P30 were measured. **(C)** Antigen-specific T cell responses. 10^5^ splenocytes from each pig were seeded on ELISpot plate and stimulated with 5 µg/mL of purified proteins or peptides. PMA/ionomycin cocktail and medium were included as positive and negative stimulation control, respectively. IFN-γ ELISpot was performed in 36 hours post stimulation. The number of SFCs per 10^5^ splenocytes was calculated and shown. P30-IRES and P30-P2A refer to P30-specific IFN-γ ELISpot following P30-IRES-MTE and P30-P2A-MTE immunization, respectively. IRES-MTE and P2A-MTE refer to MTE-specific IFN-γ ELISpot following P30-IRES-MTE and P30-P2A-MTE immunization, respectively. **(D-E)** Pearson correlations for IgG responses **(D)** and IFN-γ response **(E)** evaluated in pigs (X-axis) and mice (Y-axis). Opened circles represent individual CD2v, EP153R, P72, Penton, P54, P22, E199L, P30 antigens. Arrows in **(E)** point to MTE. Equations for the linear regression and Pearson correlation coefficients along with P values are shown. Shading area shows the filled error bars.

Antigen-specific IgG responses, as indicated by the endpoint titers, became detectable in all immunized pigs 14 days after the first immunization, increased steadily afterwards, and reached the highest level at day 35, i.e., 14 days after boost (**Figure 3B**). As observed in mice, the levels of antigen-specific IgG titers varied significantly among the different antigens. Among the 9 antigens, P72, P22, E199L and P30-IRES-MTE induced the highest titers of antigen-specific IgG responses at day 35, followed by Penton and P30-P2A-MTE, with the lowest for CD2v, EP153R, and P54. Similarly, LNP-mRNA immunization induced significant antigen-specific T cell responses as indicated by secretion of cytokines IFN-γ following re-stimulation of splenocytes with either recombinant proteins or P72 peptides or MTE peptides (**Figure 3D**). T cell responses also varied significantly among different antigens. For example, P30-IRES-MTE stimulated highest numbers of antigen-specific IFN-γ-secreting spots (103), followed by CD2v (74) and P30-P2A-MTE (73). These results show that mRNA expressing the selected ASFV antigens induces both antibody and T cell responses in pigs, but the magnitudes of the immune responses vary considerably among the different antigens.

We analyzed the correlations of antibody and T cell responses for different antigens between pigs and mice. IgG responses induced by the same antigens in pigs and mice were highly correlated with Pearson correlation coefficient of 0.93 (*p*<0.005) (**Figure 3D**). Similar, T cell-mediated IFN-γ responses induced by the same antigens in pigs and mice were also highly correlated with Pearson correlation coefficient of 0.87 (p<0.005) (**Figure 3E**). The similar results between mice and pigs cross-validate the two studies and also suggest that mice can be used to replace pigs for assessing immunogenicity of ASF subunit vaccines in most cases.

### Distinct effector functions by antibodies specific for different ASFV antigens

Due to the large size and structural complexity of virion, ASFV can infect host macrophages through multiple mechanisms, including receptor-mediated endocytosis, clathrin-mediated endocytosis, macropiniocytosis, and phagocytosis. As a result, viral neutralization assay with sera from immunization with a single ASFV antigen is not reliable (*7*) and antibody-mediated neutralization is not sufficient to confer protection (*8*). To identify potential host-dependent antiviral functions of antibodies induced by different ASFV antigens, we determined their effector functions, including ADCD, ADCC and ADCP.

To measure ADCD, CHO cells stably expressing individual ASFV antigens were incubated with heat-inactivated pig serum, followed by addition of non-heat-inactivated serum from placebo pigs as a source of complement, and cell lysis was quantified by flow cytometry (**Figure 4A** and **S4A**). ADCD activities of sera varied depending on the immunizing antigens, but overall, the sera from EP153R, P72, Penton, and P30-IRES-MTE-immunized pigs had significantly higher ADCD activities than sera from CD2v, P22, P54, E199L, and P30-P2A-MTE-immunized pigs (25-31% vs. 9-17%, **Figure 4B**).

**Fig. 4.**
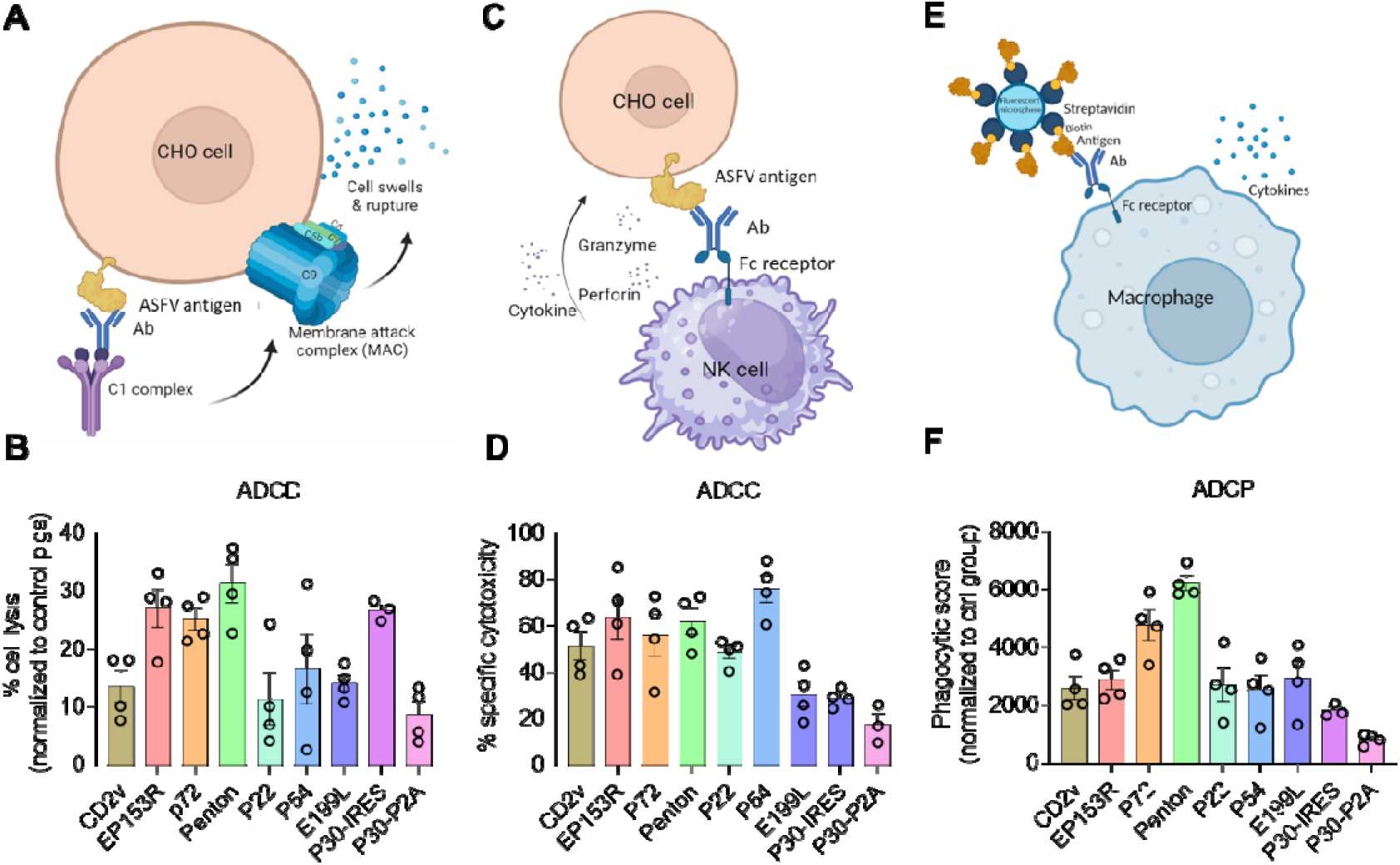
Evaluation of antibody effector functions. (**A**) Schematic diagram of ADCD assay. CHO cells stably expressing each of the selected ASFV antigens were incubated with heat-inactivated serum from immunized pigs (antibody source), followed with serum from non-immunized pigs (complement source). Lysis of CHO cells were quantified by flow cytometry. (**B**) Comparison of percentages of lysis of CHO cells among sera from pigs immunized with the indicated ASFV antigens. (**C**) Schematic diagram of ADCC assay. CHO cells stably expressing each of the selected ASFV antigens were incubated with heat-inactivated serum from immunized pigs (antibody source), followed with incubation with total PBMC from non-immunized pigs (NK cell source). Lysis of CHO cells were quantified by a luminescent assay. (**D**) Comparison of percentages of lysis of CHO cells among sera from pigs immunized with the indicated ASFV antigens. (**E**) Schematic diagram of ADCP assay. Fluorescent beads conjugated with each of the selected ASFV antigens were incubated with heat-inactivated serum from immunized pigs (antibody source), followed with incubation 3D4/31 cells. Phagocytosis of beads by 3D4/31 was quantified by flow cytometry. (**F**) Comparison of phagocytic scores among sera from pigs immunized with the indicated ASFV antigens. P30-IRES and P30-P2A refer to P30-specific antibodies following P30-IRES-MTE and P30-P2A-MTE immunization, respectively. Each circle represents one pig.

To measure ADCC, CHO cells stably expressing individual ASFV antigens were incubated with heat-inactivated pig serum, followed by addition of pig peripheral blood mononuclear cells (BPMCs) as a source of natural killer (NK) cells, and lysis of CHO cells was quantified by a luminescent assay (**Figure 4C**). Sera from P54-immunized pigs induced highest CHO cell lysis (76%), followed by sera from EP153R (64%), Penton (61.2%), P72 (56%), CD2v (51.2%), and P22 (48.6%) immunized pigs, and sera from E199L, P30-IRES-MTE and P30-P2A-MTE had lowest cell lysis (19-30%) (**Figure 4D**).

To measure ADCP, fluorescent beads were coated with specific ASFV proteins and incubated with immune sera and a pig macrophage cell line 3D4/31, and phagocytosis of labeled beads was quantified by flow cytometry (**Figure 4E** and **S4B**). Sera from P72 and Penton-immunized pigs induced highest levels of ADCP (**Figure 4F**), which were 2-3-fold higher than the ADCP induced by sera from pigs immunized with CD2v, EP153R, P22, P54 or E199L. Notably, ADCP activity was lowest for sera from pigs immunized with P30-P2A-MTE, despite robust anti-P30 IgG responses.

These results show that antibodies induced by different ASFV antigens exhibit different effector functions, which could be exploit for developing cocktail vaccines with desired immune profiles.

### Identification of antigen combinations for cocktail vaccines by computational analysis

To identify optimal antigen combinations, we performed computational analysis of five immune parameters: T cell response (IFN-γ), antigen-specific IgG level, ADCD, ADCC, and ADCP. Average values from four pigs per antigen were calculated and normalized by ranking each immune parameter from 1 (lowest) to 8 (highest). The ranking revealed distinct patterns: P22 and P30 induced the highest IgG responses; CD2v and EP153R induced the highest T cell response (IFN-γ); Penton and EP153R induced antibodies showing the highest ADCD activities; P54 and EP153R induced antibodies with the highest ADCC activities; and Penton and P72 induced antibodies with the highest ADCP activities (**Figure 5A**). When the magnitude of each immune parameters from individual antigen was taken into consideration, similar patten was also found in the chord diagram showing correlations between antigens and immunological categories (**Figure 5B**). Correlation analysis of the five immune parameters across all antigens indicated a positive correlation between Fc-mediated ADCD and ADCP, and between ADCC and T cell response (IFN-γ) (**Figure 5C**). However, IgG response was negatively correlated with IFN-γ and ADCC.

**Fig.5.**
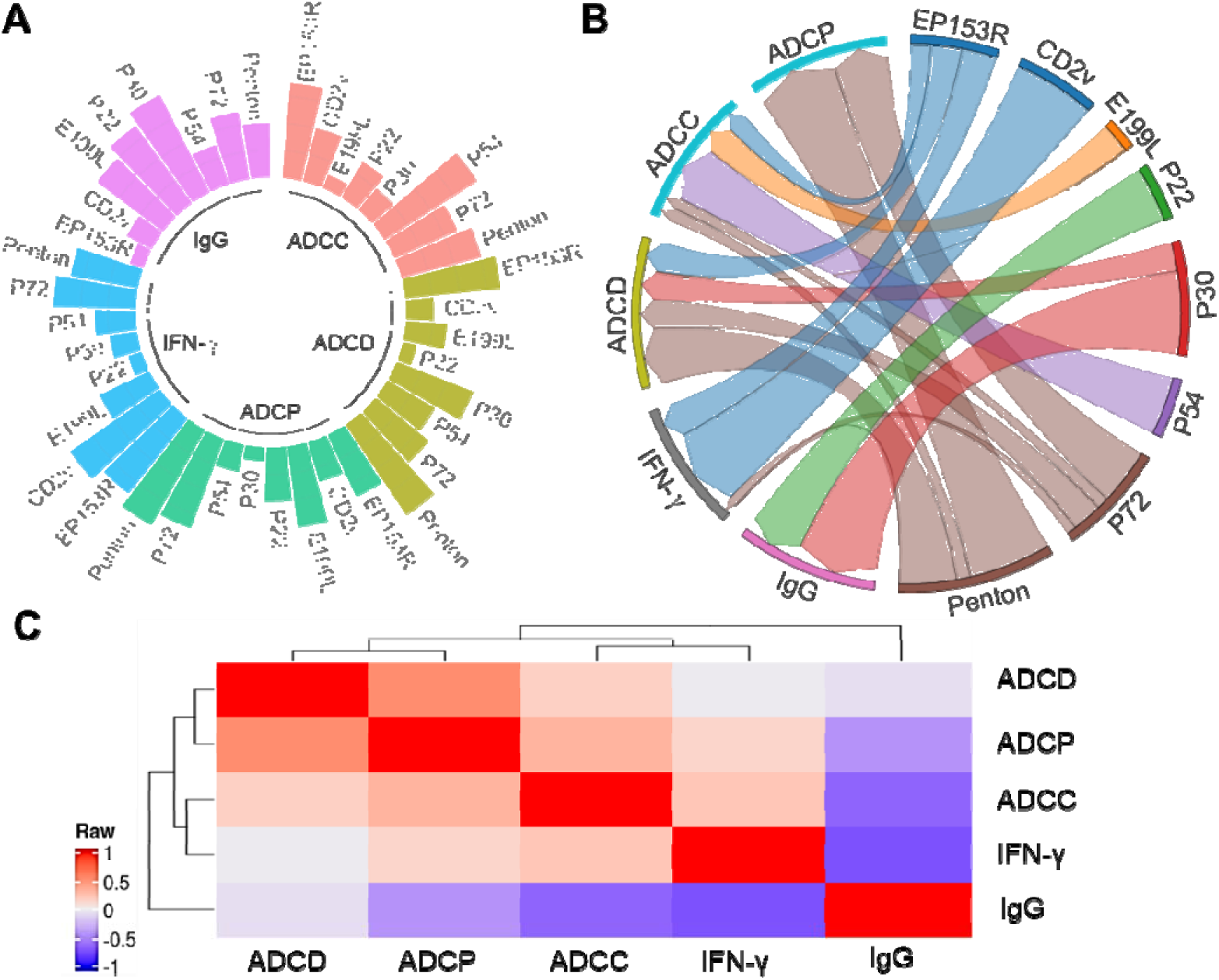
Computational analysis of immune profiles of specific antigens. (**A**) The polar bar plot of rank scores (1 to 8) of the indicated antigens in five immunological categories. The ggplot2 library in R is used to generate the Polar bar plot of rank scores of antigens. (**B**) The chord diagram of correlations between the indicated antigens and five immunological categories. Based on the average Z-scores of antigens in five immunological categories, the pyCirclize module in python3 is used to generate the chord diagram of correlations between antigens and five immunological categories, where the direction of connections is from antigens to immunological categories, and the width of the connections is proportional to the positive Z-scores. (**C**) Pearson correlation analysis of five immunological categories.

Optimal antigen combinations were scored by summing the ranks of selected immune parameters. Among the eight antigens tested (CD2v, EP153R, P72, Penton, P22, P54, E199L, P30), we developed all possible 3-way, 4-way, and 5-way combinations based on rank sums of the five immune parameters (“IgG”,“IFN-γ”,“ADCD”,“ADCC”,“ADCP”) (**Figure 6**). For the highest response across five, four or three parameters, the optimal 3-way combination was EP153R_P72_Penton (**Figure 6A**) and the optimal 4-way combination was EP153R_P54_P72_Penton (**Figure 6B**). Although the top 5-way combinations across five, four or three parameters were different, but they all included EP153R, P72 and Penton (**Figure 6C**). When ADCP was excluded because of the potential to enhance ASFV infection of macrophages due to antibody-dependent enhancement (*46, 47*), the best 3-way cocktail remained EP153R_P72_Penton, with addition of P54 for the 4-way and addition of P54 and P30 for the 5-way combinations. Additionally, when only IgG and T cell responses were considered, the top 3-way combinations were CD2v_P30_P72 or CD2v_ E199L_P72, the top 4-way combination was CD2v_E199L_P30_P72, and the top 5-way combination was CD2v_E199L_P30_P72_Penton (**Figure 6**). Although MTE-induced T cell responses and CD2v antibody-mediated hemadsorption inhibition activity were not considered here, optimal combinations for targeted immune profiles provide novel insights for developing cocktail mRNA vaccines.

**Fig. 6.**
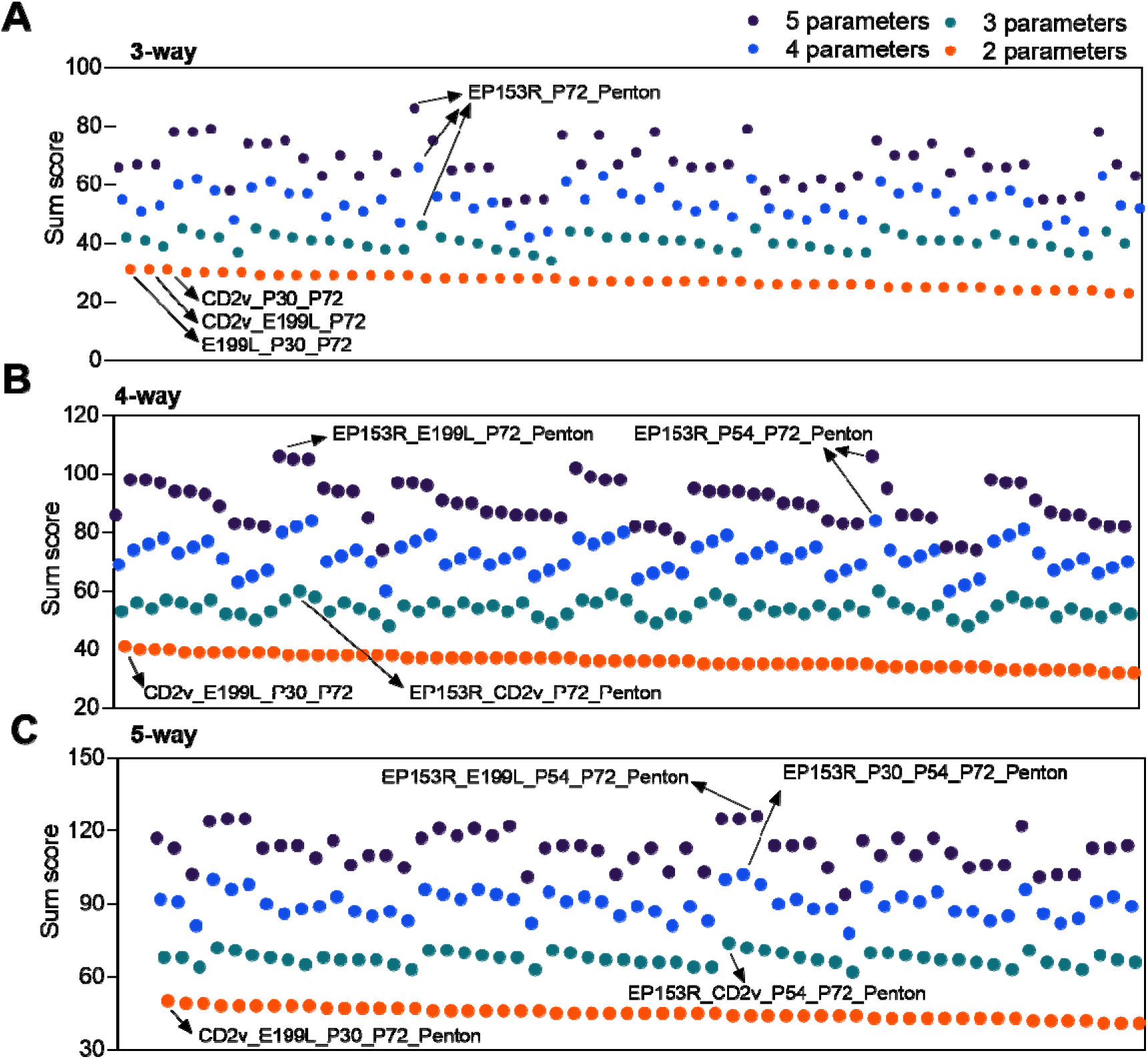
Comparison of rank scores of antigen combinations for cocktail vaccines. 3-way (**A**), 4-way (**B**), and 5-way (**C**) antigen combinations based on rank sums of 5 immune parameters (“IgG”,“IFN-γ”,“ADCD”,“ADCC”,“ADCP”), 4 parameters (“IgG”,“IFN-γ”,“ADCC”,“ADCD”), 3 parameters (“IgG”,“IFN-γ”,“ADCC”), and 2 parameters (“IgG”,“IFN-γ”) colored in black, blue, green, and red, respectively. Arrows point to the combinations with highest sum scores in different categories.

### Induction of robust antibody and T cell responses by cocktail mRNA vaccine candidates

We evaluated cocktail vaccines with antigen combinations based on computational analysis and the following considerations. All the cocktails contained P30-IRES-MTE and P72 with the former stimulating robust T cell immunity and the latter as a major capsid protein for inducing antibody responses with partial protection (**Figure 7A**). P72 was also identified as one of the three antigens in the optimal 3-way combination based on the highest response across five, four or three parameters above (**Figure 6**). CD2v was included in cocktails 1-3 to provide maximal induction of T cell immunity due to abundant T cell epitopes present in CD2v and induction of antibodies that inhibit hemadsorption, which is critical for controlling viral spread (*24, 48*). Compared to cocktail 1, penton was included in cocktail 2 as penton is a key capsid component, was identified as one of the three antigens in the optimal 3-way combination, and no study has examined whether antibodies against penton is protective. Cocktail 3 contains another highly immunogenic protein, E199L and Cocktail 4 contained P22.

**Fig. 7.**
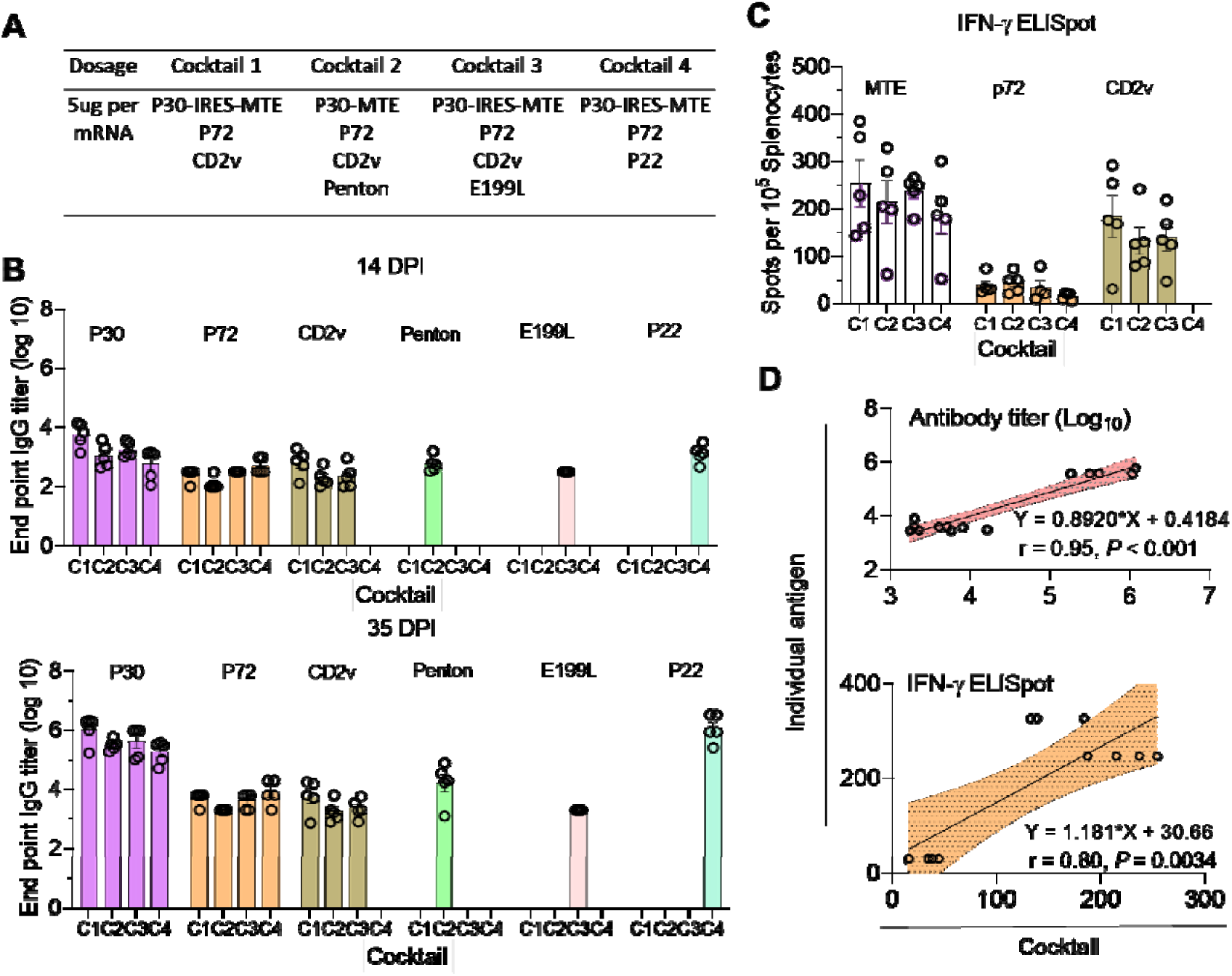
Induction of robust immune responses by candidate cocktail mRNA vaccines in mice. **(A)** Composition of the four candidate cocktail vaccines. 5 µg per mRNA was used to prepare LNP formulation. Four groups of mice (n=5 per group) were immunized at day 0 and 21. Blood samples were collected before immunization and 14 and 35 days after immunization for assaying antigen-specific IgG titers in the serum. Spleen was collected at day 35 after immunization for assaying IFN-γ secretion. **(B)** Comparison of antigen-specific IgG titers at day 14 and 35 after immunization for the indicated antigens. **(C)** Comparison of IFN-γ ELISpots in splenocytes following stimulation with MTE peptide cocktail, P72 peptides, and CD2v protein. **(D)** Pearson correlations for IgG (top) and T cell (bottom) responses induced by cocktail vaccination (X-axis) and individual antigen vaccination (Y-axis). Each circle represents one antigen: P30, P72, CD2v, Penton, E199L, P22 (top), and MTE, P72, CD2v (bottom). Equations and correlation coefficients are shown. C1, C2, C3 and C4 represent cocktail 1, cocktail 2, cocktail 3 and cocktail 4, respectively.

We evaluated the immunogenicity of candidate cocktail vaccines in mice with 5 μg of each mRNA in LNP formulations, the same amount as used in individual immunizations. BALB/c mice were immunized with the cocktail vaccines twice at day 0 and day 21 or given PBS as control. Blood was collected before immunization and at day 14 and 35 after immunization for assessing antigen-specific IgG responses by ELISA. Spleen was collected at day 35 for assessing antigen-specific T cell responses by ELISPOT. Antigen-specific IgG was detected 14 days after the first immunization and the titers were significantly increased (6-528-fold) 14 days following boost (**Figure 7B**). At day 35, IgG titers for P30 and P22 were the highest, the same as observed in mice immunized with P30-IRES-MTE or P22 individually (**Figure 2B**). The IgG titers for the rest of antigens were similar, ranging from 2×10^3^ to 1.6×10^4^. Similarly, MTE induced the highest level of T cell response as indicated by the numbers of IFN-γ ELISPOTs in all cocktails (**Figure 7C**), followed by CD2v, with the lowest for P72.

To further determine whether the immune responses elicited by specific antigens in the cocktail vaccines parallel to those elicited by individual antigens, Pearson correlation analysis was conducted. The analysis revealed correlation coefficients of 0.95 for IgG antibody responses and 0.80 for T cell responses (**Figure 6D**). Similar levels of antigen-specific IgG and T cell responses induced by specific antigens from immunization with individual LNP-mRNA versus the cocktail LNP-mRNA suggests that the inclusion of 3 or 4 mRNA in the same LNP formulation does not interfere with induction of immune responses to each antigen in the cocktail. These results also show that the cocktail ASFV vaccines induce potent antibody and T cell responses.

## Discussion

As the first step toward developing a safe and effective subunit vaccine for ASF, in this study we rationally selected and designed viral antigens, evaluated their immunogenicity in mice and pigs, developed cocktail vaccines with specific immune response profiles, and validated the induction of robust humoral and cellular immunities of the selected combinations. We chose mRNA platform for a potential ASFV subunit vaccine for the following reasons: First, although immunological correlates for a protective ASFV vaccine is not fully defined, studies have shown that antibody responses elicited by inactivated virus were not sufficient to protect pigs from infection (*8*), whereas immune responses induced by live attenuated virus confer protection (*9*). Therefore, T cell immunity likely plays a pivotal role in the protection, and an effective ASFV vaccine must induce both humoral and cellular immunities. Second, in mRNA vaccine, immunogens are expressed in native conformation by the host cells and have a higher chance to induce antibodies that recognize the native viral antigens (conformational epitopes), as well as induce CD8^+^ T cell responses that are effective in clearing virus from infected cells. Third, multiple mRNAs can be easily formulated into the same LNP, greatly simplifying the development of a cocktail vaccine.

A key aspect of subunit vaccine development is the identification of immunogens to include in the vaccine. This is especially challenging for ASFV. ASFV has a large genome, encoding more than 150 open reading frames, and many viral proteins have not been characterized. Although some viral antigens, such as P72, P30 and P54, have been shown to confer partial protection by inducing neutralizing antibodies, viral antigens that induce protective immune responses have not been defined. In addition, ASFV has complex structures and both EEV and IMV are infectious. Likely, multiple viral antigens must be combined in order to induce protective immunity.

In our selection and design of ASFV antigens, we used the following four novel approaches: First, we selected ASFV antigens that are homologs of the subunits of the protective VACV vaccine (*13, 14*). *Asfarviridae* is phylogenetically most closely related to *Poxviridae*. A VACV subunit vaccine with two antigens from EEV and two antigens from IMV confers complete protection against lethal challenge in mice (*14, 15, 49*), which was correlated with induction of serum-neutralizing antibodies and vaccinia virus-specific CD8^+^ T cells (*50, 51*). Thus, we selected the corresponding ASFV antigens based on the same localization on virion, similar functions, and amino acid sequence homology (Table 1). Second, we selected ASFV antigens based on the recent identification of P72 as the major capsid protein and Penton as a minor capsid protein (*20*). In particular, Penton has never been investigated as a vaccine antigen. Third, even for promising ASFV antigens identified previously, we designed antigen through protein engineering in order to induce strong immune responses. For example, we expressed P72 and Penton, both are intracellular proteins, in membrane-bound form, which form multimeric structures without viral chaperone and induced stronger antibody responses (*21*). Fourth, we designed T cell-directed (MTE) antigen to induce broad and robust cellular immunity. MTE contains 27 experimentally identified and computationally predicted T cell epitopes with majority being CD8 T cell epitopes. To facilitate MTE expression, we linked MTE to the abundantly expressed P30 through three strategies, GGGS linker, P2A, and IRES. *In vivo* testing of the three MTE designs showed that LNP-mRNA expressing P30-IRES-MTE stimulated strongest T cell responses (Figures 2 and 3), probably due to the direct initiation of MTE expression via IRES element. Based on these four criteria, we selected eleven ASFV antigens: two each from the outer membrane and capsid, four from the inner membrane, and one from the viral shell (Figure 1A).

We evaluated the immunogenicity of the selected ASFV antigens in LNP-mNRA formulation in both mice and pigs with concordant results. However, the magnitude of antibody and T cell responses induced by different ASFV antigens was quite different. For example, P30, P22, P72 and E199L induced robust antibody responses, but T cell responses to P30 and P22 were low (Figures 2 and 3). In contrast, CD2v and EP153R induced only modest antibody responses, probably due to their high level of glycosylation (Figure S3), but induced robust T cell responses. Our findings are in line with previous reports of abundant T cell epitopes harbored by CD2v and EP153R (*30*). Furthermore, consistent with previous observations, anti-sera induced by CD2v mRNA vaccine are effective in inhibiting hemadsorption, suggesting the functionality of the induced antibodies. Immunogenicity of E199L and Penton has not been investigated, our results show that E199L is quite immunogenic and the engineered Penton (in membrane-bound form with N180Q mutation to prevent glycosylation) also induce both antibody and T cell responses. Our results also show that MTE antigen induced the highest level of T cell responses, suggesting the validity of our design. Induction of divergent antibody and T cell responses by different ASFV antigens suggest the need to strategically combine antigens for inducing a balanced immune response for optimal protection (see below).

Another major challenge of ASFV vaccine development is a lack of reliable viral neutralizing assay due to infection of host macrophages by both EEV and IMV through receptor-mediated endocytosis, clathrin-mediated endocytosis, macropiniocytosis, and phagocytosis. This is further compounded by the requirement of BSL3 containment for any live ASFV work. To get around these obstacles, we determined effector functions, including ADCD, ADCC and ADCP, of anti-sera from pigs immunized with different ASFV antigens. The same as antibody and T cell responses, anti-sera induced by different antigens exhibited divergent effector functions (Figures 4 and 5). Nevertheless, Fc-mediated effector functions, ADCD and ADCP, were correlated, so as ADCC and T cell response (IFN-γ) (**Figure 5C**). Interestingly, IgG response was negatively correlated with IFN-γ and ADCC. To develop cocktail vaccines with desired immune response profiles, we used computational analysis to rank order antigen combinations based on the five semi-quantitative immune parameters: the levels of IgG responses, the levels of T cell responses (IFN-γ), ADCD, ADCC and ADCP. Notably, the top 3-way combination based on 5-, 4-, and 3-parameters was EP153R, P72 and Penton (Figure 6A). In the 4-parameter analysis, we removed ADCP for possible concern due to antibody-dependent enhancement (ADE), which has been observed for flavivirus infection of macrophages (*52, 53*). In the 3-parameter analysis, we further removed ADCD due to its unidentified role in disease protection. In addition, we further considered MTE-induced T cell responses and inhibition of hemadsorption by CD2v-specific antibodies. We tested four cocktail vaccines and found that antibody and T cell responses induced by specific antigen in the cocktails are highly correlated with immune responses induced by single antigen immunization (Figure 7). These results suggest that strategic combination of different antigens in LNP-mRNA vaccines are likely achieve specific immune responses profiles for optimal protection.

In summary, we have used a comprehensive approach to dissect the immune response profiles of rationally selected ASFV antigens, and developed new strategies for antigen combination for developing a safe and effective mRNA-based subunit vaccine for ASFV. The methodologies developed in this study could also be applied to develop subunit vaccines for other pathogens with large genomes, including the monkeypox virus.

### Limitation of the study

Our study has two primary limitations: First, we did not assay neutralizing activity of antibodies induced by different ASFV antigens. This is because developing neutralization assays for antibodies elicited by a single ASFV antigen has been challenging due to multiple entry pathways used by different infectious viral particles. The presence of neutralizing antibodies has been associated with live vaccine-induced protection in one report (*54*), whereas many studies suggest that neutralizing antibodies to P30, P54, and P72 are insufficient for protection (*8*), highlighting the critical role of T cell responses in vaccine-induced protection (*55*). Given the multilayered structures of ASFV and its various entry mechanisms, both neutralizing antibodies and cellular immunity are likely required for blocking viral infection and clearing infected cells. Additionally, antibodies can control infection through mechanisms beyond neutralization. Due to the limited studies on ASFV, the roles of Fc-mediated complement recruitment and engagement of innate immune cells, such as NK cells and macrophages, should not be underestimated, as previous reports have shown that antibody effector functions are associated with protection against respiratory syncytial virus (*56*), influenza virus (*57*), and malaria parasites (*58*). Our study delineates the antibody effector functions and other immune parameters by selected ASFV antigens, offering critical insights for the development of cocktail vaccines. Second, we did not assess protective efficacy of cocktail vaccines under challenge conditions due to the select agent status, restricted biocontainment requirements, and high cost for working with ASFV infection in pigs. Clearly, this should be a priority for moving forward. In addition, we selected ASFV antigens based on the current knowledge of ASFV structure and protein functions. It cannot be excluded that other proteins exhibiting similar localization or functions may be discovered in the future and are worthwhile for testing.

## Materials and Methods

### Cells and plasmids

Human embryonic kidney cells (HEK-293T) and Vero E6 cells were maintained in minimum essential medium supplemented with 10% fetal bovine serum, antibiotics (100 units/ml of penicillin and 100 mg/ml of streptomycin) and fungizone (0.25 mg/ml) at 37°C with 5% CO_2_. Porcine macrophage 3D4/31 cell line was maintained in RPMI with essential supplements including FBS, antibiotics, and fungizone. FreeStyle™ CHO-S cells were obtained from ThermoFisher and cultured in Freestyle™ CHO Expression Medium supplemented with 8 mM L-glutamine at 37℃, 8% CO_2_ on an orbital shaker platform rotating at 135 rpm. ASFV P72 (B646L), Penton (H240R), CD2v (EP402R), P22 (KP177R), EP153R (C-type lectin), E248R, E199L, P54 (E183L), P30 (CP204L) genes from Georgia 2007/1 strain (NCBI Reference Sequence: NC_044959.2) were synthesized by GenScript and inserted into the phCMV mammalian cell expression vector (MoBiTec). P72 and Penton were engineered for membrane anchoring by addition of secretion signal peptide from human CD8α (GenBank ID: NP_001139345.1) to the N terminus and CD8α stalk region or hinge, transmembrane region, and a short cytoplasmic tail to the C terminus (*21*).

### Design of T cell-directed vaccine

ASFV-derived CD4^+^ and CD8^+^ T cell epitopes were searched on the Immune Epitope Database (IEDB) (https://www.iedb.org/) and limited to those with positive experimental results for IFN-γ ELISpot and/or cellular MHC/mass spectrometry ligand presentation. A total of 22 T cell epitopes from PP62, MGF100-1L, A238L, K145R, MGF505-7R, P34, P150, P37, and EP153R were selected for their strong induction of cellular immunity according to previous publications (*10, 30, 37, 40, 59, 60*). P72 and CD2v contains multiple T cell epitopes which were not included in the T cell-directed vaccine design. Additionally, the online server, NetMHCpan 4.1 (http://www.cbs.dtu.dk/services/NetMHCpan/) was used to predict binding affinities of all 8-11mer peptides in the most abundantly expressed pp220 polyprotein. The most frequent Swine Leukocyte Antigens (SLA-1:0401, SLA-1:0101, SLA-1:0801) were selected for peptide binding. Strong binders with an %Rank-Eluted ligand (EL) <0.1 and affinity <200 mM was selected and included 3 epitopes from M448R and 2 epitopes from MGF505-7R. The T cell-directed vaccine construct, multiple-T cell epitope (MTE), contained a Kozak sequence followed by P30, Internal Ribosome Entry Site (IRES) or porcine teschovirus-1 2A (P2A) or GGGS linker, and individual T cell epitopes with additional 5 amino acid residues (AA) on each side were fused together, plus a HA tag for MTE detection. MTE was synthesized by GenScript and cloned into expression vector with P30, namely P30-MTE, P30-IRES-MTE, and P30-P2A-MTE.

### Lipid Nanoparticle (LNP) formulation of mRNA and validation

Genes encoding the eleven candidate antigens were inserted into pUC57 vector, which contains essential elements for mRNA *in vitro* transcription, such as T7 promoter, UTRs, and polyA tail. The linearized plasmids were subject to in vitro transcription using pseudouridine-5’-triphosphate or pseudo-UTP (*44*). After mRNA purification and capping reaction, an aqueous phase of mRNA was prepared by diluting mRNA stock in 10 mM citrate buffer. The organic phase of lipid nanoparticles was prepared by adding 200 proof ethanol with lipid stock solutions which contained ionizable lipid L002 (Advanced RNA Vaccine, ARV), DSPC helper lipid (Avanti Polar Lipids), cholesterol (Avanti Polar Lipids), and DMG-PEG-2000 (Avanti Polar Lipids). LNP-mRNA formulations were prepared by mixing organic and aqueous phases at a ratio of 1:3. RiboGreen assay (Thermo Fisher) was performed by following manufacture’s instruction to quantify mRNA after formulation. Encapsulation efficiency was measured by Picogreen assay. To validate LNP delivery of mRNAs, HEK-293T cells were seeded onto 24-well tissue culture plate and 500 ng of LNP-mRNA diluted in Opti-MEM was added in individual well. Cells were collected in 48 h and protein expression was analyzed by flow cytometry and confocal microcopy as described below.

### Flow cytometry

HEK-293T cells seeded in 6-well tissue culture plate and transfected with 3 µg recombinant phCMV plasmids expressing individual proteins using the linear 25 kDa polyethylenimine (PEI; Santa Cruz Biotechnology) at a 3:1 mass ratio of PEI to DNA. At 48 h post transfection, cells were trypsinized and washed with 1 mL FACS buffer followed by direct surface staining of P72 using mouse anti-ASFV P72 mAb (MyBioSource, San Diego, CA), mouse anti-P30 monoclonal antibody (Aviva Systems Biology, OAEF00154), P54 mAb (GeneTex, GTX635690), or mouse mAb specific for HA tag (GenScript). Alexa Fluor™ 488-conjugated Goat anti-Mouse IgG (H+L) was used as the secondary antibody and cells nuclei were counterstained with DAPI (4’,6-Diamidino-2-Phenylindole, Dilactate) before data acquisition on a BD LSR Fortessa HTS-2 cytometer. Results were analyzed using FlowJo v10 software.

### Confocal microscopy

Vero E6 cells were grown on glass-bottom 35-mm cell culture dishes (MatTek). In 48 hours after transfection of MTE expression vectors, cells were fixed by 4% paraformaldehyde at room temperature (RT) for 15 min followed by permeabilization with 0.5% Triton X-100 for 10 min and then blocked with 2% bovine serum albumin for 30 min. Cells were incubated with mouse anti-P30 mAb (GenScript) and Rabbit anti-HA tag polyclonal antibody (Thermo Fisher) at 37C for 1 hour. Alexa Fluor™ 488-conjugated goat anti-mouse IgG and Alexa Fluor® 594 AffiniPure™ goat anti-rabbit IgG (H+L) (Jackson ImmunoResearch) were used as secondary antibody. Cells were counterstained with DAPI before proceeding with imaging under a confocal microscope (Nikon A1R HD25, Nikon). Images were processed using the program ImageJ (https://imagej.net/Fiji).

### Mouse study

To assess immunogenicity of individual antigens in mice, six to eight-week-old female BALB/c mice were purchased from Charles River and housed in animal facility at the Massachusetts Institute of Technology (MIT). All animal ethical and welfare standards were met in this study and experiments were approved by the Institutional Animal Care and Use Committee at MIT under protocol number 0322-021-25. Briefly, 5 mice were assigned to each group and immunized intramuscularly with 50 µL 5 µg LNP-mRNA diluted in PBS. For assessing immunogenicity of candidate vaccine cocktails, a total of 50 µL composed of 5 µg of each selected antigens were injected to the hind limb of a mouse. All mice were boosted three weeks later. Serum samples were collected from submandibular vein prior to immunization and two weeks after each injection. Mice were euthanized at day 35 and spleen tissue was collected for analysis of cellular immunity.

### Pig study

LNP-mRNA vaccination of pigs was performed according to the protocols approved by Committee on Animal care (protocol number 2308000566) and Midwest Veterinary Service (MVS), Inc. (protocol number 24005). A total of 38 four-week-old piglets were randomly assigned into ten groups with four pigs for each of the 9 LNP-mRNAs groups (MB-P72, MB-Penton, CD2v, P22, E199L, EP153R, P54, P30-IRES-MTE, and P30-P2A-MTE) and the rest two pigs assigned to control group injected with sterile PBS. 30 ug of LNP-mRNA expressing each individual antigen was diluted to 1 mL in sterile PBS and injected to the back of ear intramuscularly and boosted three weeks later. Serum samples were collected before vaccination and weekly after each injection. Body weight of each pig was measured before the study and at the termination. All pigs were euthanized two weeks after boost and spleen, draining dorsal superficial cervical (DSCLN) and mandibular (MLN) lymph nodes were collected for testing T cell responses.

### Indirect ELISA

Recombinant ASFV proteins with a HIS tag were inserted into PET28a vector and expressed in a BL21 *Escherichia coli* system under induction of isopropyl-β-D-thiogalactopyranoside (IPTG), followed by protein purification via Ni-nitrilotriacetic acid (NTA) agarose. For detection of antibody response in serum from immunized animals, 300 ng purified proteins were coated onto 2HB ELISA plate (Thermo Fisher) in antigen coating buffer (35 mM sodium bicarbonate and 15 mM sodium carbonate, pH 8.8) and incubated at 37C overnight (*41*). Plates were washed with PBST (0.05% Tween 20 in 1× phosphate-buffered saline) three times and blocked with 5% non-fat milk in PBST for 2 hours at room temperature. 100 µL two-fold serial dilutions of serum starting from 1:50 were added to the coated plate and incubated for 2 hours at room temperature. HRP-conjugated goat anti-mouse or pig IgG (H+L) (Thermo Fisher) was used as the secondary antibody. Colorimetric reaction was developed by ABTS peroxidase substrate system and stopped by ABTS stop solution in 30 min after color development. optical density OD_405_ value was obtained to quantify the coloring intensity using the Tecan Infinite® 200 PRO microplate reader. Cutoff values for each protein were determined by OD value of control group animal samples plus three standard deviations. The endpoint titer for each animal that has the highest dilution giving a reading above cutoff was calculated by interpolating from a sigmoidal standard curve using GraphPad Prism.

### Enzyme-linked immunospot (ELISPOT) assay

To evaluate T cell immunity, 10^5^ splenocytes from each mouse or pig were cultured in CTL medium and seeded on pre-coated mouse IFN-γ/TNF-α Double-Color ELISPOT plate (Cellular Technology Limited) or IFN-γ single-color ELISpot plate. Cells from each animal were stimulated with 5 µg/mL P72 peptide cocktail (**Table S2**), MTE peptide cocktail, or purified proteins (CD2v, P22, EP153R, Penton, P54, P30). Cocktail of phorbol 12-myristate 13-acetate (PMA) and ionomycin (Thermo Scientific) was used as positive control and cell culture medium as negative control. Plate was incubated at 37 °C with 5% CO_2_ for 36 hours followed by washing with distilled H_2_O for three times before addition of primary and secondary antibodies based on manufacture’s protocol (Cellular Technology Limited). Colored spots were counted by an automated immunospot analyzer (Cellular Technology Limited). The antigen-specific IFN-γ and TNF-α spot-forming cells (SFCs) were calculated and results were analyzed using GraphPad Prism.

### Hemadsorption inhibition assay

CD2v has been reported to contribute to hemadsorption which is a term describing ASFV-infected cells attracting red blood cells to attach on cell surface and form “rosette” pattern (*45*). To establish an assay for measuring anti-CD2v antibody-mediated hemadsorption inhibition, HEK 293T cells transfected with CD2v-expression vector were harvested and mixed with 1:100 diluted serum pretreated with Receptor Destroying Enzyme (RDE) at 37℃ for 18h and then 56℃ for 1h. After 1 hour incubation of serum with cell, 2% red blood cells (RBC) were added and incubated at 37℃ for 24 hours. Monolayer cells were gently washed with PBS for five times and fixed, permeabilized, and blocked for antibody staining. Rabbit anti-RBC polyclonal antibody (LSBio) and mouse anti-HA mAb (Thermo Fisher) were added and incubated for 1 hour at 37℃. Alexa Fluor™ 488-conjugated goat anti-mouse IgG and Alexa Fluor® 594 AffiniPure™ goat anti-rabbit IgG (H+L) (Jackson ImmunoResearch) were used as secondary antibody. Cells were counterstained with DAPI before proceeding with imaging by confocal microscopy (Nikon A1R HD25, Nikon). Images were processed using the program ImageJ (https://imagej.net/Fiji) and geometric mean fluorescent intensity of AF594 red fluorescence for RBC were calculated and compared between groups.

### Antibody-dependent complement deposition (ADCD) assay

Before testing, pig serum samples were inactivated at 56℃ for 30 min. CHO-S cells stably expressing each of the ASFV proteins (CD2v, EP153R, P22, P54, E199L and membrane-bound P72, P30, Penton) were used as target cells and seeded in non-tissue culture treated 96-well plates at a density of 1.0 × 10^5^ cells per well. 10-fold diluted inactivated serum were added and mixed briefly with the cells. After incubation for 30 min at 37℃ in a humidified 5% CO_2_ incubator, 50 µL control pig serum was added into each well as the complement source and plates were incubated at 37C for 18 hours, followed by DAPI staining. Cells were subject to flow cytometry using a BD LSR Fortessa HTS-2 cytometer. Results were analyzed using FlowJo v10 software and the percentage of live CHO cells positive for mCherry (%live) were calculated. The complement-mediated cytotoxicity was calculated according to the formula: cytotoxicity (%) = ((1 – (%live of sample / %live of no serum ctrl)) × 100. The antigen-specific cytotoxicity effect was calculated by normalizing to the control group serum.

### Antibody-dependent cellular cytotoxicity (ADCC) assay

PBMCs collected from control pigs were used as a source of NK cells and pre-activated with IL-2 (final concentration 20ng/mL) and IL-12 (final concentration 25 ng/mL) at 37℃ for 24 hours. CHO cells expressing ASFV proteins were seeded on 96-well plates at a density of 2 × 10^4^ cells per well. Inactivated serum was 1:10 diluted and incubated with CHO cells for 30 min at 37℃. 50 µL of 2% Triton was added as positive control for complete cell lysis. 4 × 10^5^ Pre-activated PBMCs were added to achieve an effector to target cell ratio of 20:1. Following 24 hours incubation at 37℃ in humidified incubator, a CytoTox-Glo™ luciferase based assay was used to measure the luminescence of viable and non-viable cells according to manufacturer’s instructions. Percentage of cell lysis (% lysis) was calculated by dead cell luminescence divided by total cell luminescence. The percentage of antigen-specific cytotoxicity was calculated using the formular: (% lysis of testing sample - % lysis of no serum ctrl) / (% lysis of triton - % lysis of no serum ctrl).

### Antibody-dependent cellular phagocytosis (ADCP) assay

ASFV proteins were biotinylated using EZ-Link™ Micro Sulfo-NHS-LC-Biotinylation Kit and then conjugated to 1 µm red FluoSpheres™ NeutrAvidin™-Labeled Microspheres by overnight incubation at 4℃. Fluorescent microspheres were centrifuged at 13,000 g for 2 min at 4℃ and washed twice with 1% BSA in PBS to remove the excess unbound proteins. The antigen-coated microspheres were resuspended in 1% BSA. Effector cells were porcine macrophage 3D4/31 cell line labeled with CellTrace™ CFSE Cell Proliferation Kit. To perform the phagocytosis assay, CFSE-labeled 3D4/31 cells were seeded on 96-well plate at a density of 1 × 10^4^ cells per well and incubated for 24 hours in 5% CO_2_ incubator. Cell culture media were removed and treated with fucoidan (final concentration 100 µg/mL) for 1 hour at 37℃ to block scavenger receptors. Inactivated serum was 10-fold diluted and mixed with 10 μL suspension (equivalent to 4 × 10^6^ antigen-coated microspheres). After 30 min incubation at 37 ℃, the mixture was transferred to fucoidan-treated cells and incubated for 6 hours. Cells were washed extensively with sterile PBS, trypsinized, and resuspended in 200 µL medium containing DAPI. Cell suspension was subject to flow cytometry. Phagocytosed microspheres were gated as double positives for CFSE-labeled cells and fluorescent beads. Phagocytic score was determined by multiplying the percentage of fluorescent-bead-positive cells by the mean fluorescent intensity (MFI) of this population. The antigen-specific phagocytosis was computed by normalizing the phagocytic score to control group pigs.

### Statistical analysis

Data was processed using R version 4.4.1 (*61*) with tidyverse version 2.0.0 (*62*). The R environment was executed using singularity version 3.5.0 (*63*) and this docker container docker://bumproo/bulk_r441. Four biological replicates for each antigen, except for P30 which had three replicates, were summarized by averaging. Correlation analysis was done using the cor function from the stats package in R with the pearson method and heatmaps were plotted using the Heatmap function from the R package ComplexHeatmap version 2.20.0 (*64*). These averaged values were then normalized by ranking antigen values for each immunological category from 1 (lowest) to 8 (highest). The ggplot2 library in R is used to generate the Polar bar plot of rank scores of antigens based on the average rank scores. Based on the average Z-scores of antigens in five immunological categories, the pyCirclize module in python3 is used to generate the chord diagram of correlations between antigens and five immunological categories. All possible 5-way, 4-way, and 3-way antigen combinations were then scored by summing the ranks. The sum scores calculated for the following sets of immunological data: (“IgG”,“IFN-γ”,“ADCD”,“ADCC”,“ADCP”), (“IgG”,“IFN-γ”,“ADCC”,“ADCD”), (“IgG”,“IFN-γ”,“ADCC”), and (“IgG”,“IFN-γ”).

All the histogram charts were expressed as means +/- standard error of mean (SEM). Differences between groups were evaluated by one-way analysis of variance (ANOVA) using Prism software version 6 (GraphPad Software). A *P* value of <0.05 was regarded as statistically significant difference.

## Acknowledgments

We thank the Koch Institute Robert A. Swanson (1969) Biotechnology Center for technical support, specifically Flow Cytometry, Microscopy, Nanotechnology Materials and Biopolymers & Proteomics, Bioinformatics Facilities.

## Author contributions

Conceptualization: FY, J Cui, J Chen

Methodology: FY, J Cui, TW, JQ, JHJ, HD, CAW

Investigation: FY, J Cui, J Chen

Supervision: J Chen, RX, HC

Writing—original draft: FY

Writing—review & editing: FY, J Chen

