## Supplemental data for "Selection, Design and Immunogenicity Studies of ASFV Antigens for Subunit mRNA Cocktail Vaccines with Specific Immune Response Profiles"

Fangfeng Yuan *et al.*

**This PDF file includes:**

Supplementary Text

Figs. S1 to S4

Tables S1 to S2

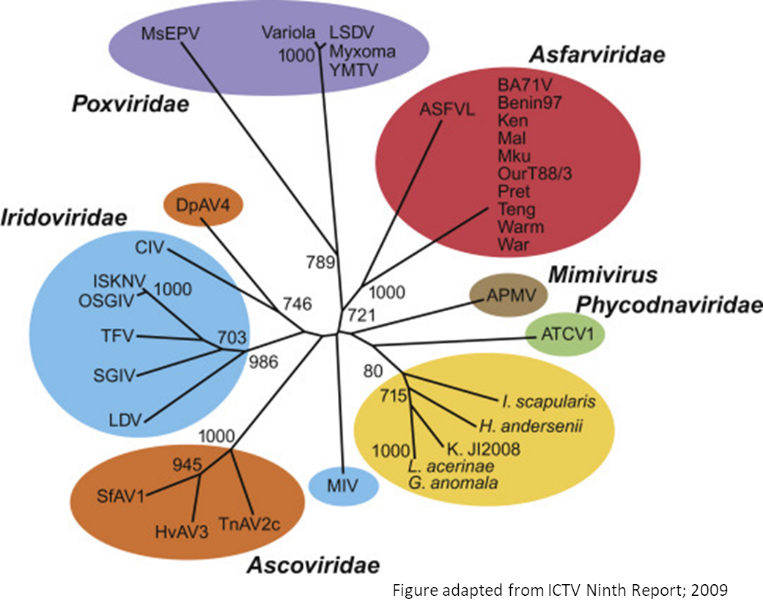
Fig. S1. Phylogeny of *Nucleocytoviricota* phylum. Figure is adapted from International Committee on Taxonomy of Viruses (ICTV) Ninth Report, 2009.

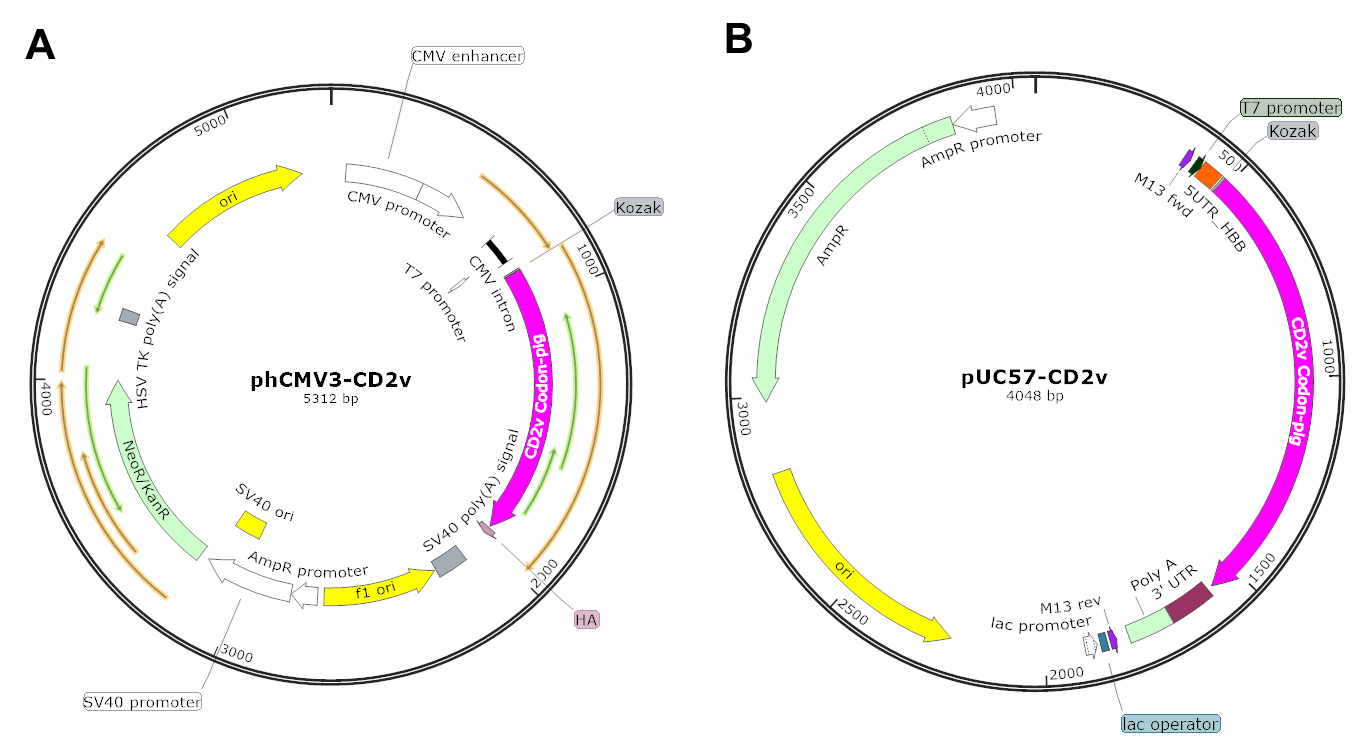
Figure S2: Representative plasmid maps for CD2v protein expression (A) and mRNA synthesis (B). DNA constructs encoding ASFV genes were shown as an example with detailed elements.

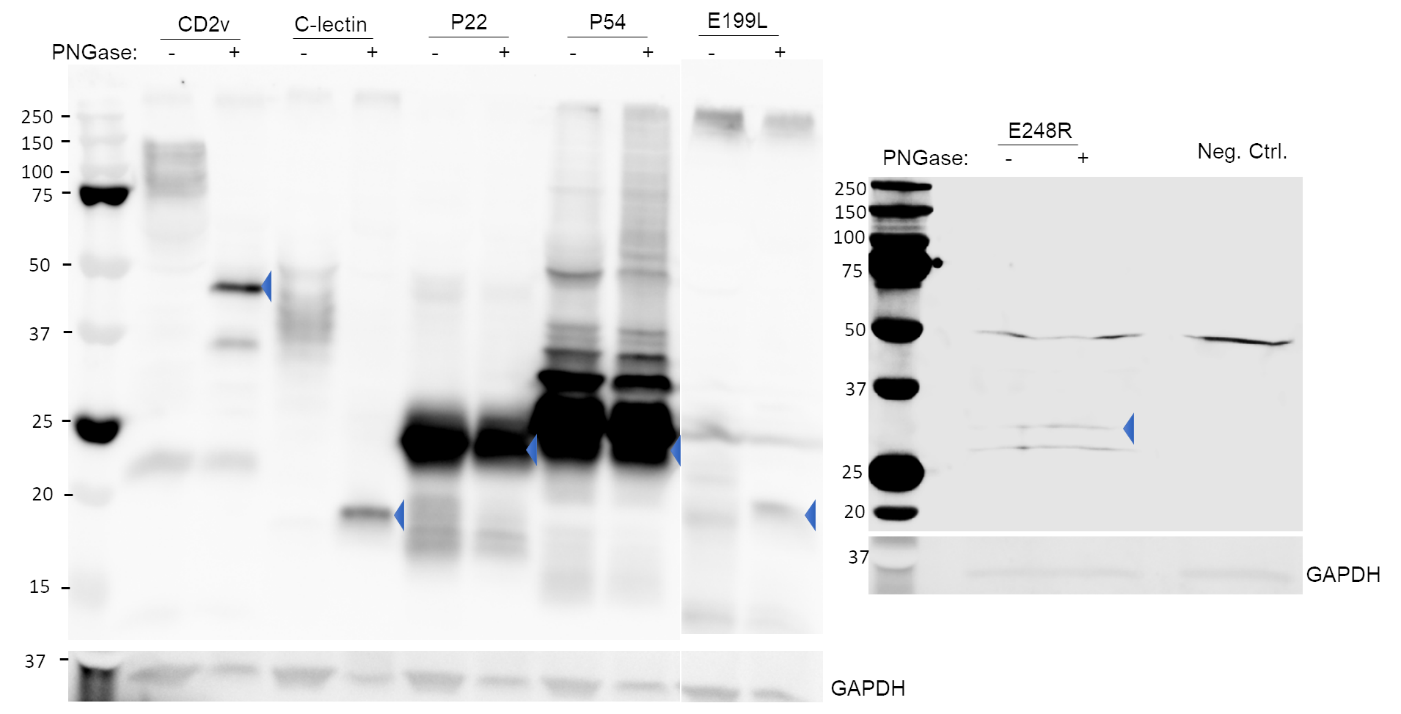
**Figure S3: Validation of antigen expression by Western blotting**

HEK-293T cells were transfected with plasmids expressing individual HA-tagged ASFV antigens and cell lysate was collected for Western blotting at 48 hours post transfection. Cell lysates were not treated or treated with PNGase to remove the N-linked glycans before SDS-PAGE. Membranes were probed with a rabbit polyclonal antibody against HA tag, followed by a StarBright™ Blue 700 conjugated goat anti-rabbit IgG. The signals were visualized by a Bio-Rad ChemiDoc MP Imaging System. Blue triangles point to the predicted size of ASFV antigens.

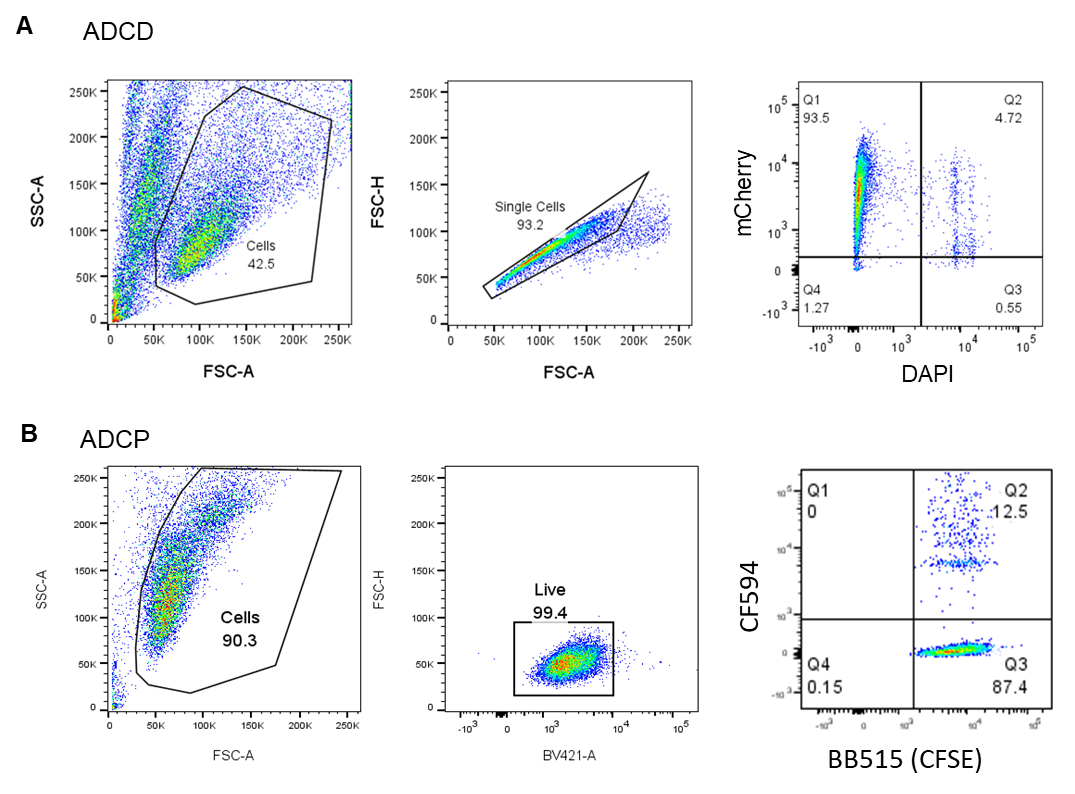
Fig. S4. Representative gating strategies for ADCD and ADCP assays. To measure complement-mediated cell lysis (A), following incubation with heat-inactivated test serum and non-heat-inactivated pig serum as a complement source, CHO cells stably expressing individual ASFV antigens plus mCherry were resuspended in FACS buffer containing DAPI, followed by flow cytometry. Single cells were gated and plotted for DAPI versus mCherry. Live CHO cells in Q1 were identified as mCherry+ DAPI- and dead CHO cells in Q2 were identified as mCherry+ DAPI+. Percentages of ADCD-mediated cell lysis were calculated using the indicated formula in the methods. To measure ADCP (B), fluorescent beads (CF594) were conjugated with specific ASFV proteins and incubated with immune sera and CFSE-labeled pig macrophage cell line 3D4/31, and phagocytosis of labeled beads was quantified by flow cytometry as positive for both CFSE and CF594. Flow plots show FSC vs. SSC (left), live cells (middle), and CFSE vs. CF594 (right).

Table S1. Physicochemical properties LNP-mRNAs

| Formulation | Hydrodynamic size(nm) | Polydispersity index | Encapsulation efficiency (%) |
| --- | --- | --- | --- |
| LNP-mRNA-E199L | 89.53 ± 0.11 | 0.116 ± 0.01 | 89.6 |
| LNP-mRNA-P72 | 85.31 ± 0.11 | 0.101355 ± 0.03 | 86.2 |
| LNP-mRNA-IRESMTE | 94.265 ± 0.15 | 0.114045 ± 0.02 | 84.3 |
| LNP-mRNA-P2AMTE | 88.55 ± 0.31 | 0.13485 ± 0.04 | 88.5 |
| LNP-mRNA-EP153R | 87.25 ± 0.86 | 0.12375 ± 0.01 | 81.9 |
| LNP-mRNA-P24 | 101.75 ± 0.21 | 0.1167 ± 0.03 | 82.4 |
| LNP-mRNA-P54 | 105 ± 1.84 | 0.1383 ± 0.05 | 86.7 |
| LNP-mRNA-CD2v | 106.65 ± 2.33 | 0.083835 ± 0.02 | 80.7 |
| LNP-mRNA-Penton | 102.85 ± 0.21 | 0.17535 ± 0.02 | 84.8 |

Table S2. Peptide pool used for inducing P72- and MTE-specific T cell responses in vitro

| **Name** | **Peptide Sequence** |
| --- | --- |
| P72 | SSYIPFHYGGNAIK |
|  | SRISNIKNVNKSY |
|  | SLDEYSSDVTTL |
| MTE | SYYTKAENI |
|  | RYVKDVLPL |
|  | AFINSTDFL |
|  | VPTRLHSFL |
|  | NYWVNYSLI |
